# GBP1 recruitment to actin-rich pedestals induced by extracellular Gram-negative bacteria promotes pyroptosis

**DOI:** 10.1101/2025.09.25.678451

**Authors:** Daniel J Bennison, Ishaan Chaudhary, Dharitri Chaudhuri, Justin Chun Ngai Wong, Ayush Punwatkar, Priyanka Biswas, Miyu Stephenson, Qiyun Zhong, Wouter W Kallemeijn, Marianne Guenot, Sandra Koigi, Diana Papp, John P Thomas, Dimple Dixit, Tamas Korcsmaros, Arthur M Talman, Eva-Maria Frickel, Edward W Tate, Sandhya S Visweswariah, Gad Frankel, Avinash R Shenoy

## Abstract

The IFNγ-induced GTPase guanylate binding protein 1 (GBP1) binds to lipopolysaccharide (LPS) on cytosolic Gram-negative bacteria and promotes pyroptosis via the recruitment and activation of caspase-4 on the bacterial outer membrane. Enteropathogenic and enterohaemorrhagic *Escherichia coli* (EPEC and EHEC, respectively) are extracellular pathogens that also induce LPS-and caspase-4-dependent pyroptosis. However, whether GBP1 is involved in this process remains unknown. EPEC and EHEC adhere intimately to intestinal epithelial cells via avid interactions between the bacterial adhesin Intimin and Tir (Translocated intimin receptor), a type 3 secretion system effector protein. Intimin-mediated clustering of Tir triggers actin polymerisation, leading to pedestal-like structures at bacterial attachment sites. Here we show that GBP1 is recruited to actin-rich pedestals in human cells infected with EPEC and EHEC *in vitro* and mouse colonocytes infected with the EPEC-like murine pathogen *Citrobacter rodentium in vivo*. GBP1-dependent caspase-4 trafficking to these sites leads to pyroptosis and IL-18 release. GBP1 mutants defective in LPS coatomer formation also localised to EPEC pedestals, pointing to LPS-independent mobilisation of GBP1 by actin polymerisation induced by extracellular A/E pathogens. To further dissect the underlying mechanism, we engineered a chimeric receptor (FcγR-Tir) by combining the intracellular signalling domain of Tir and the extracellular ligand-binding domain of the Fcγ receptor. Clustering of FcγR-Tir with IgG-coated beads produced ‘sterile’ actin-rich pedestals that were sufficient to recruit GBP1 independently of bacteria. Our findings reveal that cytosolic GBP1 is mobilised to sites of pathogen-induced actin remodelling independently of LPS. We establish that GBP1 not only operates as a pattern-recognition receptor but also orchestrates effector-triggered immunity against pathogens that hijack the actin cytoskeleton.

**175-word Abstract:** The IFNγ-induced GTPase guanylate binding protein 1 (GBP1) binds to lipopolysaccharide (LPS) on cytosolic Gram-negative bacteria and promotes pyroptosis via the recruitment and activation of caspase-4 on the bacterial outer membrane. Enteropathogenic and enterohaemorrhagic *Escherichia coli* (EPEC and EHEC, respectively) are extracellular pathogens that adhere to host cells and stimulate dense actin polymerisation underneath their attachment sites, generating structures called actin-rich ‘pedestals’. Here we show that GBP1 traffics to actin-rich pedestals in human cells infected with EPEC and EHEC *in vitro* and mouse colonocytes infected with the EPEC-like murine pathogen *Citrobacter rodentium in vivo*. GBP1 promotes caspase-4 recruitment to actin-rich pedestals, leading to pyroptosis and IL-18 release. GBP1 mutants defective in LPS coatomer formation also localised to EPEC pedestals. Our novel assay that mimics pathogenic effector activity revealed GBP1 recruitment to ‘sterile’ actin polymerisation sites. We conclude that cytosolic GBP1 is mobilised to sites of pathogen-induced actin remodelling independently of LPS. Our study establishes that GBP1 not only operates as a pattern-recognition receptor but also orchestrates effector-triggered immunity against pathogens that hijack the actin cytoskeleton.

## Introduction

Guanylate Binding Protein 1 (GBP1) is the forerunner of a family of seven human IFNγ-inducible 65-75 kDa GTPases that promote antimicrobial responses and inflammasome activation in response to intracellular pathogens (<u>Kim *et al*, 2011</u>; <u>Kutsch & Coers, 2021</u>; <u>Rivera-Cuevas *et al*, 2023</u>; <u>Shenoy *et al*, 2012</u>; <u>Tretina *et al*, 2019</u>). GBP expression can be induced by IFNγ in most cell types, reflecting their broad roles in cell-autonomous immunity (<u>Tretina *et al*., 2019</u>). GBPs were first implicated in pyroptosis via mouse caspase-4 (also called caspase-11) in macrophages exposed to cytosolic LPS (<u>Finethy *et al*, 2017</u>; <u>Meunier *et al*, 2014</u>; <u>Pilla *et al*, 2014</u>). More recently, we and others have shown that human GBP1 drives pyroptosis by enabling the recruitment and activation of caspase-4 on the surface of cytosolic Gram-negative bacteria (<u>Fisch *et al*, 2019</u>; <u>Kutsch *et al*, 2020</u>; <u>Santos *et al*, 2020</u>; <u>Wandel *et al*, 2020</u>). GBP1 directly binds LPS and forms polymeric ‘coats’ on the outer membrane of bacteria (e.g., *Shigella flexneri, Burkholderia thailandensis, Francisella novicida*, *Salmonella enterica* serovar Typhimurium, *Yersinia*) (<u>Kirkby *et al*, 2023</u>; <u>Kuhm *et al*, 2025</u>; <u>Kutsch *et al*., 2020</u>; <u>Weismehl *et al*, 2024</u>; <u>Zhu *et al*, 2024</u>). GBP1 is proposed to act as a surfactant that extracts LPS and facilitates its binding to caspase-4, thereby stimulating its proteolytic activity (<u>Kutsch *et al*., 2020</u>; <u>Shi *et al*, 2014</u>). Active caspase-4 cleaves gasdermin D (GSDMD) leading to pyroptosis, and generates the bioactive form of IL-18, which promotes intestinal homeostasis and antimicrobial immunity (<u>Shi *et al*, 2023</u>). In macrophages, caspase-4 activation leads to downstream ‘non-canonical’ activation of the NLRP3 inflammasome leading to caspase-1-dependent maturation of IL-1β (<u>Kayagaki *et al*, 2011</u>). GBP1 thus defends against invasive cytosolic Gram-negative bacteria by acting as a pattern-recognition receptor (PRR) for LPS to promote the pyroptotic removal of bacterial niches and release of immune cytokines.

Biochemically, the GTPase activity of GBP1, which drives GBP1 dimerisation and higher-order oligomerisation, is required for its recruitment to purified LPS and outer membrane of intact Gram-negative bacteria (<u>Kutsch & Coers, 2021</u>; <u>Rivera-Cuevas *et al*., 2023</u>; <u>Tretina *et al*., 2019</u>). In addition, GBP1 has a C-terminal farnesylation motif (^589^CTIS^592^) and a proximal polybasic region (^584^RRRK^587^), which contribute to GBP1 trafficking to cytosolic bacteria and pyroptotic responses (<u>Kohler *et al*, 2020</u>; <u>Piro *et al*, 2017</u>). Other GBP family members, such as GBP2-5, may support pyroptosis, but their involvement is pathogen- and cell type-dependent (<u>Fisch *et al*., 2019</u>; <u>Kutsch *et al*., 2020</u>; <u>Santos *et al*., 2020</u>; <u>Wandel *et al*., 2020</u>). In addition, GBP1 targets other intracellular bacteria (e.g., *Listeria*, *Mycobacterium*), protozoan parasites (e.g., *Toxoplasma gondii*) and viruses (e.g. HIV) (<u>Kutsch & Coers, 2021</u>), suggesting that GBP1 may detect signals other than LPS. The role of GBP1, if any, in detecting extracellular bacterial infection remains unexplored.

Enteropathogenic and enterohaemorrhagic *E. coli* (EPEC and EHEC, respectively) exemplify extracellular Gram-negative human pathogens that trigger caspase-4-dependent pyroptosis (<u>Sanchez-Garrido *et al*, 2020</u>). EPEC is among the leading causes of childhood mortality and morbidity in low- and middle-income countries (<u>Chen & Frankel, 2005</u>; <u>Kotlof f *et al*, 2013</u>), and EHEC causes fatal infection mainly in high-income countries (<u>Launders *et al*, 2016</u>). These pathogens tightly attach to gut epithelia, manipulate the host actin cytoskeleton, and trigger the loss of brush border microvilli leading to characteristic attaching and effacing (A/E) lesions (<u>Frankel *et al*, 1998</u>). EPEC and EHEC remain on the outside of the host cell and deploy a type 3 secretion system (T3SS) to inject 20-50 effectors to manipulate host signalling (<u>Shenoy *et al*, 2018</u>). Clustering of the T3SS effector Tir on the host plasma membrane by the bacterial outer membrane adhesin Intimin triggers actin polymerisation leading to the formation of structures commonly referred to as bacterial ‘pedestals’ (<u>Frankel & Phillips, 2008</u>). We previously showed that actin polymerisation by Tir^EPEC^ or Tir^EHEC^ drives rapid LPS- and caspase-4-dependent pyroptosis of human macrophages (<u>Goddard *et al*, 2019</u>). Moreover, during infection of IFNγ-primed non-phagocytic intestinal epithelial cells by EPEC, bacterial LPS can be internalised into cells in a Tir-dependent manner, leading to caspase-4 activation and pyroptosis (<u>Zhong *et al*, 2022</u>; <u>Zhong *et al*, 2020</u>).

The Tir proteins of EPEC and EHEC have a hairpin topology and an Intimin-binding region within the extracytoplasmic loop. However, their C-terminal regions use distinct mechanisms to polymerise actin. Intimin-mediated clustering of Tir^EPEC^ results in its tyrosine phosphorylation (on Y474 in Tir^EPEC^) by Src tyrosine kinases, thereby generating a binding site for the adaptor proteins NCK1/2 (<u>Kenny *et al*, 1997</u>; <u>Phillips *et al*, 2004</u>). These adaptors recruit the actin nucleation-promoting factor N-WASP (Neuronal Wiskott-Aldrich syndrome protein), which stimulates ARP2/3-dependent actin polymerisation (<u>Gruenheid *et al*, 2001</u>). In contrast, Tir^EHEC^ recruits N-WASP-ARP2/3 via a second T3SS effector, the Tir-cytoskeleton coupling protein (TccP; also called *E. coli* secreted protein F-like from prophage U (EspF_U_)) (<u>Campellone *et al*, 2004</u>; <u>Garmendia *et al*, 2004</u>). The Tir^EHEC^ and TccP interaction is scaffolded by the host proteins IRTKS (Insulin receptor tyrosine kinase substrate) or IRSp53 (Insulin Receptor Substrate p53), which bind Tir^EHEC^ at its ^456^NPY^458^ motif (<u>Crepin *et al*, 2010</u>; <u>Vingadassalom *et al*, 2009</u>; <u>Weiss *et al*, 2009</u>). *Citrobacter rodentium* (CR), an EPEC-like natural murine pathogen, is commonly used to model infection *in vivo* (<u>Collins *et al*, 2014</u>; <u>Deng *et al*, 2003</u>; <u>Mullineaux-Sanders *et al*, 2019</u>). Tir mutants of CR unable to polymerise actin are less virulent *in vivo* during mouse infection, pointing to the crucial role of Tir in actin polymerisation and pathogenesis (<u>Crepin *et al*, 2015</u>).

Here we investigated the role of GBP1 in pyroptosis in response to adherent Gram-negative bacteria in non-phagocytic epithelial cells. We reveal that GBP1 is recruited to actin-rich pedestals generated by Tir-Intimin signalling by EPEC and EHEC, and that GBP1 recruitment is independent of bacterial LPS. GBP1-dependent caspase-4 recruitment to pedestals is also independent of LPS, which is only required for stimulating caspase-4 activity. Thus, pathogen-induced actin polymerisation is a novel signal that mobilises cytosolic GBP1 for effector-triggered immunity (ETI) against extracellular pathogens.

## Results

### GBP1 promotes pyroptosis during EPEC and EHEC infection

We previously showed that EPEC O127:H6 (strain E2348/69) induces caspase-4-dependent pyroptosis in IFNγ-primed colonic epithelial cells (<u>Zhong *et al*., 2020</u>). Similarly, in IFNγ-primed HeLa cells, wild-type (WT) EPEC induced pyroptosis as quantified by propidium iodide-uptake assays (***Figure 1A***) in a caspase-4- and gasdermin D (GSDMD)-dependent manner (***Figure EV1A***). In contrast, pyroptosis was not induced following infection by an isogenic T3SS-defective Δ*escF* mutant (***Figure 1A***), indicating that host cell pyroptosis requires bacterial effector translocation. Deletion of GBP1 (*GBP1^-/-^* HeLa cells) led to a marked reduction in pyroptotic lysis and IL-18 release upon EPEC infection (***Figure 1A, EV1B***). Likewise, infection with WT EHEC O157:H7 (strain 85-170), but not an isogenic T3SS-defective Δ*escN* mutant, triggered pyroptosis in a GBP1-dependent manner (***Figure 1B***). Together, these results indicate that GBP1, caspase-4, and a functional bacterial T3SS are required for pyroptosis and IL-18 release during infection by EPEC and EHEC.

**Figure 1.**
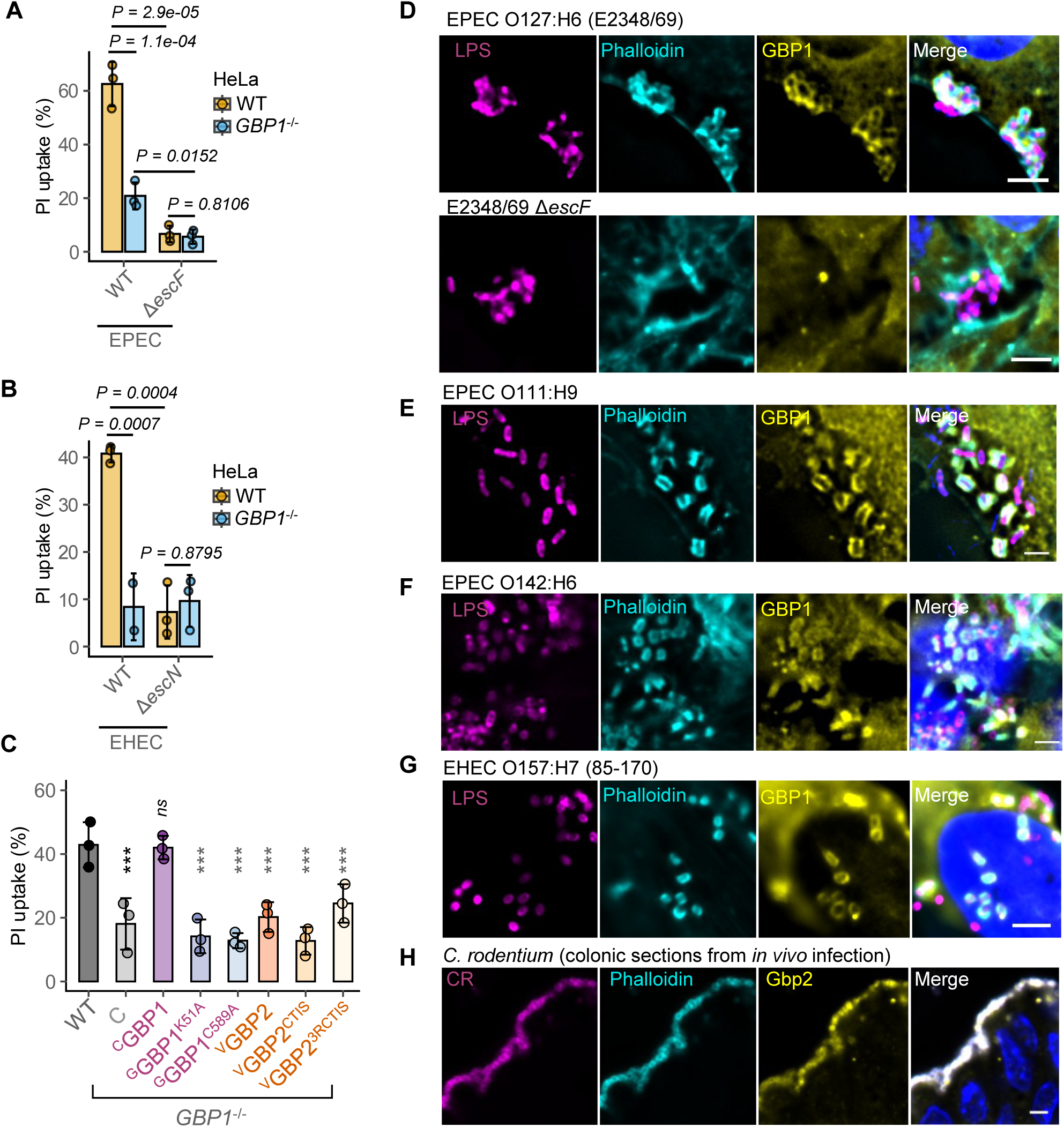
GBP1 promotes pyroptosis and is recruited to actin-rich pedestals of extracellular Gram-negative pathogens. (A) – (B) Percentage pyroptotic cell death as measured by propidium iodide dye uptake assays from IFNγ-primed wild-type (WT) or *GBP1*^-/-^ HeLa cells infected with wild-type (WT) EPEC O127:H6 (strain E2348/69) or an isogenic Δ*escF* mutant for 6 h (A) or WT EHEC O157:H7 (strain 85-170) or an isogenic Δ*escN* mutant for 8 h. Mean ± SD error bars with symbols representing data from n = 3 independent experiments are shown. * *P < 0.05, *** P < 0.001, ns – not significant (P > 0.05)* are two-tailed *P* values for the indicated comparisons from mixed effects ANOVA. (C) Percentage pyroptotic cell death as measured by propidium iodide dye uptake assays from IFNγ- and doxycycline-treated wild-type (WT) or *GBP1*^-/-^ HeLa cells stably expressing tetracycline (Tet)-inducible mCherry2 (C), mCherry2-tagged wild-type GBP1 or the K51A or C589A mutants, or mVenus (V)-GBP2 or the indicated mutants. Mean ± SD error bars with symbols representing data from n = 3 independent experiments are shown. *\*\*\* P < 0.001, ns – not significant (P > 0.05)* are two-tailed *P* values for comparisons between WT cells and cells expressing the indicated GBP1 or GBP2 variants from mixed effects ANOVA. Exact *P* values as follows – C: 2.3e-06; cGBP1: 0.7787; ^G^GBP1^K51A^: 5.3e-07; ^G^GBP1^C589A^: 4.5e-07; ^V^GBP2: 5.1e-07; ^V^GBP2^CTIS^: 4.5e-07; ^V^GBP2^3RCTIS^: 4.1e-05. (D) Representative immunofluorescence images of IFNγ-primed HeLa cells infected with WT EPEC or an Δ*escF* mutant for 2 h, as indicated. Cells were stained with anti-LPS (to stain bacteria) and anti-GBP1 antibodies, phalloidin (to stain actin) and Hoechst (DNA dye; blue) as labelled. Scale bar, 5 μm. (E) – (G) Representative immunofluorescence images of IFNγ-primed HeLa cells infected with EPEC O111:H9 (E), EPEC O142:H6 (F) or EHEC O157:H7 (strain 85-170) (G) as labelled. Infections were carried out for 3 h (E-F) or 5 h (G). Cells were stained with anti-LPS (to stain bacteria) and anti-GBP1 antibodies, phalloidin (to stain actin) and Hoechst (DNA dye; blue) as labelled. Scale bar, 5 μm. (H) Representative immunofluorescence images of colonic sections from C57BL/6 mice infected with CR at 8-day post-infection. Sections were stained with anti-CR (to stain bacteria) and anti-GBP1 antibodies, phalloidin (to stain actin) and Hoechst (DNA dye; blue) as labelled. Data are representative of images from 5 infected mice. Data from the following number (n) of independent repeats: D, n = 6; E-F, n = 2; G-H, n = 5.

GBP1 prenylation and GTPase activity are essential for its pyroptotic functions during cytosolic bacterial infection (<u>Fisch *et al*., 2019</u>; <u>Kutsch *et al*., 2020</u>; <u>Santos *et al*., 2020</u>; <u>Wandel *et al*., 2020</u>) (***Figure EV1C***). We therefore used tetracycline (Tet)-inducible expression of mCherry2-GBP1 (^C^GBP1) variants to examine GBP1 activities required for EPEC-induced pyroptosis as described before (<u>Fisch *et al*., 2019</u>; <u>Fisch *et al*, 2020</u>; <u>Fisch *et al*, 2023</u>). Expression of ^C^GBP1, but not mCherry2 (C) alone, fully complemented the defect in pyroptosis and IL-18 production in *GBP1*^-/-^cells (***Figure 1C; EV1C-E***), attesting that the loss-of-function phenotype of these cells is specifically due to *GBP1,* and mCherry2-tagging does not affect protein function. Further, the expression of the GTP binding-deficient K51A and prenylation-null C589A GBP1 variants failed to rescue pyroptosis or IL-18 release (***Figure 1C; EV1C-E***), indicating that both properties are required for pyroptosis during infection by EPEC.

Overexpression of GBP2, which naturally undergoes geranylgeranylation and lacks a polybasic motif, can restore caspase-4-dependent pyroptosis in *Shigella flexneri-*infected *GBP1*^-/-^ epithelial cells, suggesting that GBP2 may compensate for the loss of GBP1 expression (<u>Dickinson *et al*, 2023</u>). However, in contrast to *Shigella* infection, elevated expression of mVenus-GBP2 (^V^GBP2) in the *GBP1^-/-^* cells did not restore pyroptosis or IL-18 release during EPEC infection (***Figure 1C; EV1C-E***), indicating an essential role for GBP1 in this context. We also tested two chimeric variants of GBP2: one incorporating the GBP1 CTIS motif (leading to GBP2 farnesylation; ^V^GBP2^CTIS^) and another incorporating the CTIS motif plus the polybasic region of GBP1, yielding ^V^GBP2-^583^RRRKACTIS^591^ (^V^GBP2^R3CTIS^). Neither variant could restore pyroptosis or IL-18 secretion defects in *GBP1^-/-^* cells upon EPEC infection (***Figure 1C; EV1C-E***).

Altogether, GBP1 catalytic activity and prenylation are required for pyroptosis and IL-18 production in response to EPEC infection and these actions cannot be compensated by overexpression of GBP2 or with variants substituted with C-terminal regions of GBP1.

### GBP1 is recruited to actin-rich pedestals induced by extracellular bacteria

As GBP1 forms coats on the LPS of cytosolic bacteria, we used immunofluorescence microscopy to assess GBP1 localisation during infection by EPEC and related A/E pathogens. During infection, EPEC uses its bundle forming pilus (BFP) to form microcolonies on host cells (<u>Shenoy *et al*., 2018</u>). Actin-rich pedestals were formed by 100 % of microcolonies of WT EPEC at 2 h post-infection, and around 80 % of these were positive for endogenous GBP1 (***Figure 1D, EV2A***), indicating extensive recruitment of GBP1 to EPEC attachment sites. In contrast, GBP1 did not traffic to attachment sites of the Δ*escF* strain, which forms microcolonies and attaches to similar extent as WT EPEC, but cannot inject effectors or elicit actin polymerisation (***Figure 1D, EV2A***). CL40 human colonic epithelial cells exemplify well-differentiated primary-like cells with low epithelial to mesenchymal transition (EMT) signature and have an intact IFNγ response (<u>Berg *et al*, 2017</u>; <u>Chisanga *et al*, 2016</u>; <u>Roumeliotis *et al*, 2017</u>). Endogenous GBP1 in CL40 cells also colocalised with actin-rich pedestals of WT, but not Δ*escF*, EPEC (***Figure EV2B***), indicating that GBP1 localisation to EPEC pedestals also occurs in physiologically relevant intestinal epithelial cells. We also tested the response of HT29 human intestinal epithelial cells which undergo IFNγ-dependent pyroptosis during *Salmonella* infection (<u>Santos *et al*., 2020</u>). Like in HeLa and CL40 cells, GBP1 recruitment to EPEC pedestals was observed in HT29 cells (***Figure EV2C***). EPEC-induced cell death in IFNγ-primed CL40 and HT29 was markedly reduced upon stable silencing of caspase-4 and GSDMD (***Figure EV2D-E***), indicating that the pyroptotic response to EPEC infection of HeLa cells mimics intestinal epithelial cells. We next used 2D monolayers prepared from human colonic organoids to assess GBP1 trafficking in untransformed primary intestinal epithelial cells. GBP1 trafficked to the actin-rich pedestals of WT, but not Δ*escF*, EPEC in organoid monolayers, further confirming GBP1 trafficking to EPEC pedestals in physiologically relevant systems (***Figure EV2F***).

To rule out LPS O-antigen-specific recruitment of GBP1, we tested clinical EPEC isolates expressing distinct O-antigens. Actin-rich pedestals of EPEC O111:H9, O142:H6, and O119:H6, were also positive for GBP1 (***Figure 1E-F, EV2G***). This indicates that GBP1 targets EPEC-induced actin-rich pedestals in epithelial cells in a strain- and O-antigen-independent manner. Likewise, GBP1 was recruited to actin-rich pedestals of EHEC O157:H7 (***Figure 1G***). Actin-rich pedestals were formed by ∼65 % of EHEC bacteria at 5 h post-infection, with 100 % of these pedestals also positive for GBP1 (***Figure EV2H***), pointing to similarly ubiquitous targeting of EHEC pedestals. Having observed GBP1 recruitment to EPEC and EHEC pedestals *in vitro*, we asked whether GBP1 targets A/E pathogens *in vivo.* To address this, we stained sections from colons of C57BL/6 mice infected with CR at the peak of infection (8 days post-infection). We observed that Gbp2, the murine orthologue of human GBP1 (<u>Kutsch & Coers, 2021</u>; <u>Rivera-Cuevas *et al*., 2023</u>; <u>Tretina *et al.*, 2019</u>), colocalised with the actin-rich pedestals of CR (***Figure 1H***).

As GBP2-5 can also target cytosolic bacteria, we assessed whether additional GBPs targeted EPEC attachment sites. Tet-inducible FLAG- or MYC-tagged GBP2-5 did not target EPEC pedestals (***Figure EV3A***), indicating that A/E bacterial actin-rich pedestals mainly attract GBP1. GBP2 appeared to be in the vicinity of a fraction of pedestals (∼20 %), pointing to its infrequent trafficking to EPEC attachment sites (***Figure EV3A***). We therefore investigated whether GBP2 contributed to EPEC-induced pyroptosis by stably silencing its expression in HeLa and HT29 cells using our miRNA30E-based system. GBP2 expression was markedly reduced as seen by western blotting (***Figure EV3B-C***), however, EPEC-induced pyroptosis in *GBP2-*silenced HeLa or HT29 cells was similar in non-targeting control cells (***Figure EV3B-C***). In contrast, stable silencing of *GBP1* in HT29 and CL40 cells reduced EPEC-induced pyroptosis, further confirming that GBP1 promotes pyroptosis in infected intestinal epithelial cells (***Figure EV3C-D***).

GBP1 can be secreted from cells, including during bacterial infection (<u>Naschberger *et al*, 2017</u>). It was therefore plausible that GBP1 can bind to EPEC extracellularly following its secretion. To test this, we performed immunofluorescence analyses with and without permeabilisation to limit anti-GBP1 antibody access to extracellular GBP1, if any. All EPEC bacteria expressing mVenus fluorescent protein were stained with anti-LPS antibodies with or without permeabilisation, confirming that EPEC remains outside cells during infection as expected (***Figure EV3E***), and validating the experimental approach. No GBP1 colocalisation was observed with EPEC when cells were not permeabilised prior to immunostaining, revealing that GBP1 is intracellular and recruited to actin-rich structures underneath bacterial attachment sites (***Figure EV3E***). Altogether, we conclude that cytosolic GBP1 traffics to actin-rich pedestals of extracellular, intimately adherent A/E pathogens *in vitro* and *in vivo*.

### GBP1 promotes caspase-4 recruitment to bacterial pedestals

GBP1 facilitates caspase-4 recruitment to intracellular Gram-negative bacteria (<u>Fisch *et al*., 2019</u>; <u>Kutsch *et al*., 2020</u>; <u>Santos *et al*., 2020</u>; <u>Wandel *et al*., 2020</u>), therefore we tested caspase-4 localisation during A/E pathogen infection. Immunofluorescence microscopy revealed that endogenous caspase-4 localised to actin-rich pedestals of EPEC and EHEC (***Figure EV3F-G***). As both caspase-4 and GBP1 antibodies that reliably detect their respective targets were raised in mice (***Reagents and Tools Table***), we exploited the YFP-Caspase-4^C258S^ reporter (a catalytically inactive variant that serves as a fluorescent tracer without causing cytotoxicity) that we previously developed to monitor caspase-4 recruitment to cytosolic *Salmonella* (<u>Fisch *et al*., 2019</u>). Like native caspase-4, YFP-caspase-4^C258S^ localised to > 95 % of GBP1-positive pedestals of EPEC and EHEC, verifying that the YFP-caspase-4^C258S^ reporter mimics the localisation of the native protein (***Figure 2A, 2C***).

**Figure 2.**
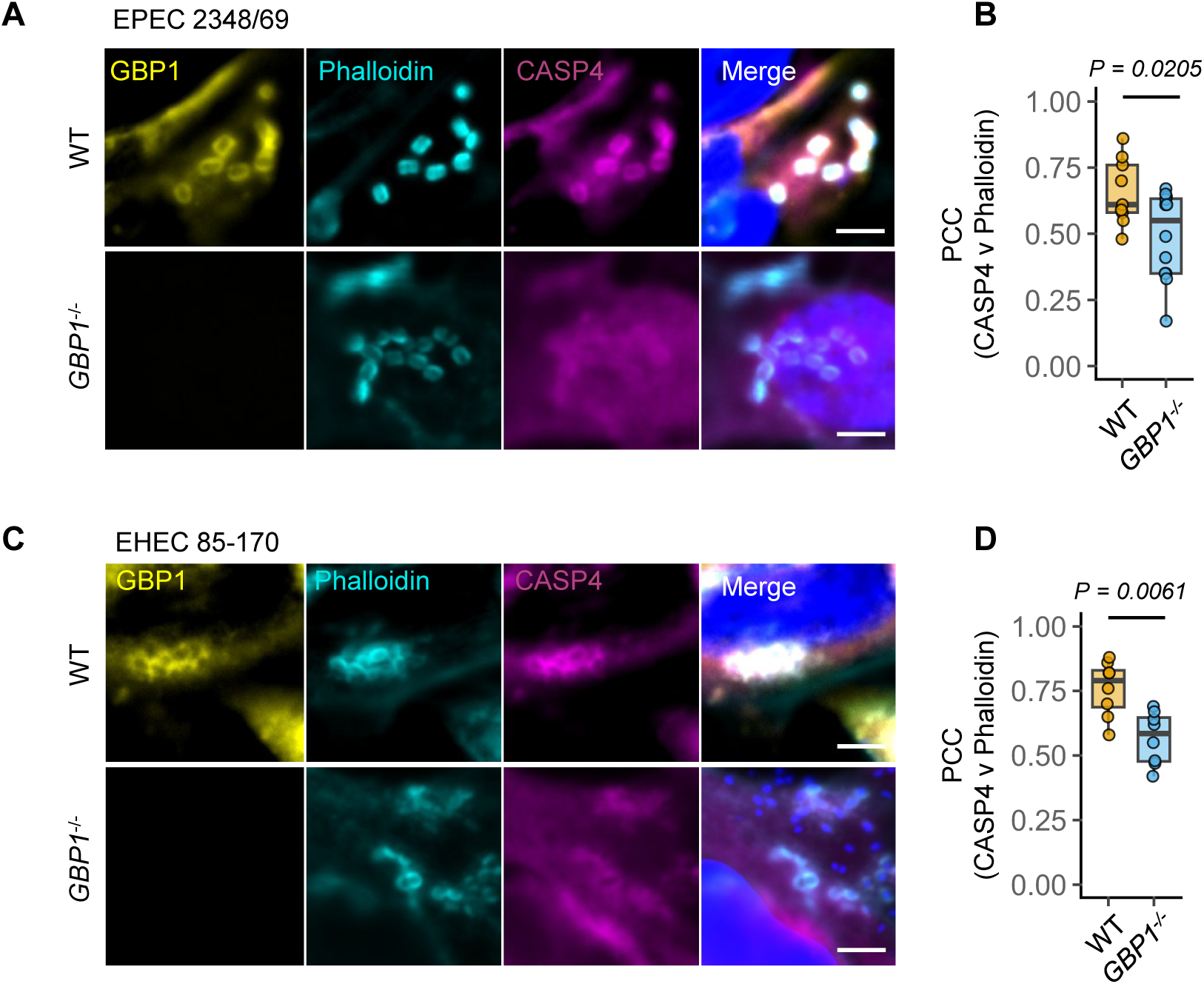
GBP1 enables caspase-4 recruitment to bacterial attachment sites. (A) Representative immunofluorescence images of IFNγ-primed wild-type (WT) or *GBP1*^-/-^HeLa cells expressing YFP-Caspase4^C285S^ infected with EPEC for 2 h. Cells were stained with anti-GBP1 antibodies, phalloidin (to stain actin) and Hoechst (DNA dye; blue). Scale bar, 5 μm. (B) Pearson’s correlation coefficient (PCC) for images from experiments in (A) assessing colocalization of caspase-4 and phalloidin are shown in WT or *GBP1*^-/-^ cells as labelled. (C) Representative immunofluorescence images of IFNγ-primed WT or *GBP1*^-/-^ HeLa cells expressing YFP-Caspase4^C285S^ infected with EHEC for 5 h. Cells were stained with anti-GBP1 antibodies, phalloidin (to stain actin) and Hoechst (DNA dye; blue). Scale bar, 5 μm. (D) Pearson’s correlation coefficient (PCC) for images from experiments in (C) assessing colocalization of caspase-4 and phalloidin are shown in WT or *GBP1*^-/-^ cells as labelled. Data from n = 5 independent experiments, and each dot in B and D represent independent fields of view used for PCC measurements across experiments are shown. Box plots, median and interquartile range (IQR) are shown, with boxes indicating IQR, whiskers indicating 1.5x IQR and the line indicating the median. Two-tailed *P* values for the indicated comparisons from an independent sample Student’s *t* test are indicated in B and D.

EPEC and EHEC adhered to similar extents and induced comparable actin pedestal formation in WT and *GBP1*^-/-^ cells, ruling out a role for GBP1 in bacterial attachment or pedestal formation (***Figure 2A, 2C***). We next queried whether caspase-4 localisation to pedestals is GBP1-dependent. Caspase-4 recruitment was reduced in *GBP1*^-/-^ cells, displaying diffused localisation throughout the cell (***Figure 2A, 2C***). In further support for GBP1-dependent recruitment of caspase-4, the Pearson’s correlation coefficient (PCC) for phalloidin (which stains polymerised actin) and caspase-4 at EPEC and EHEC pedestals in *GBP1*^-/-^ cells was significantly reduced as compared to WT cells (***Figure 2B, 2D***). Together, these results show that GBP1 is important for caspase-4 enrichment at sites of actin-polymerisation induced by these bacteria.

### GBP1 recruitment to pedestals requires Tir-driven actin polymerisation

We next investigated the role of actin polymerisation in pyroptosis and the recruitment of GBP1-caspase-4 to A/E pathogen attachment sites. EPEC mutants that cannot form actin-rich pedestals, such as the Δ*escF*, Δ*tir* and Δ*eae* (lacking the gene that encodes intimin; ***Figure EV4A***) strains failed to stimulate pyroptosis or IL-18 release (***Figure 3A-B***). Accordingly, neither GBP1 nor caspase-4 was recruited to the attachment sites of these EPEC mutants (***Figure 3C-F***). Comparable EHEC Δ*escN*, Δ*tir* and Δ*eae* mutants also did not trigger pyroptosis (***Figure EV4B***), form pedestals, or recruit GBP1 or caspase-4 (***Figure EV4C-F***). To establish whether Tir clustering, rather than ‘downstream’ actin polymerisation, is sufficient for GBP1 and caspase-4 recruitment, we used an EHEC Δ*tccP* mutant strain, which has intact Tir and Intimin proteins and can therefore lead to Tir clustering, but lacks the TccP protein to recruit N-WASP-ARP2/3 for actin polymerisation (***Figure EV4A***). As expected, EHEC Δ*tccP* bacteria did not form pedestals (***Figure EV4G***), failed to induce pyroptosis (***Figure EV4B***) and did not recruit GBP1 or caspase-4 (***Figure EV4G***). These results underscore the requirement for EPEC- or EHEC-induced actin cytoskeleton manipulation via Tir-Intimin signalling for GBP1 mobilisation, which then promotes caspase-4 recruitment to these sites. Furthermore, the recruitment of both proteins correlates with pyroptosis and IL-18 release.

**Figure 3.**
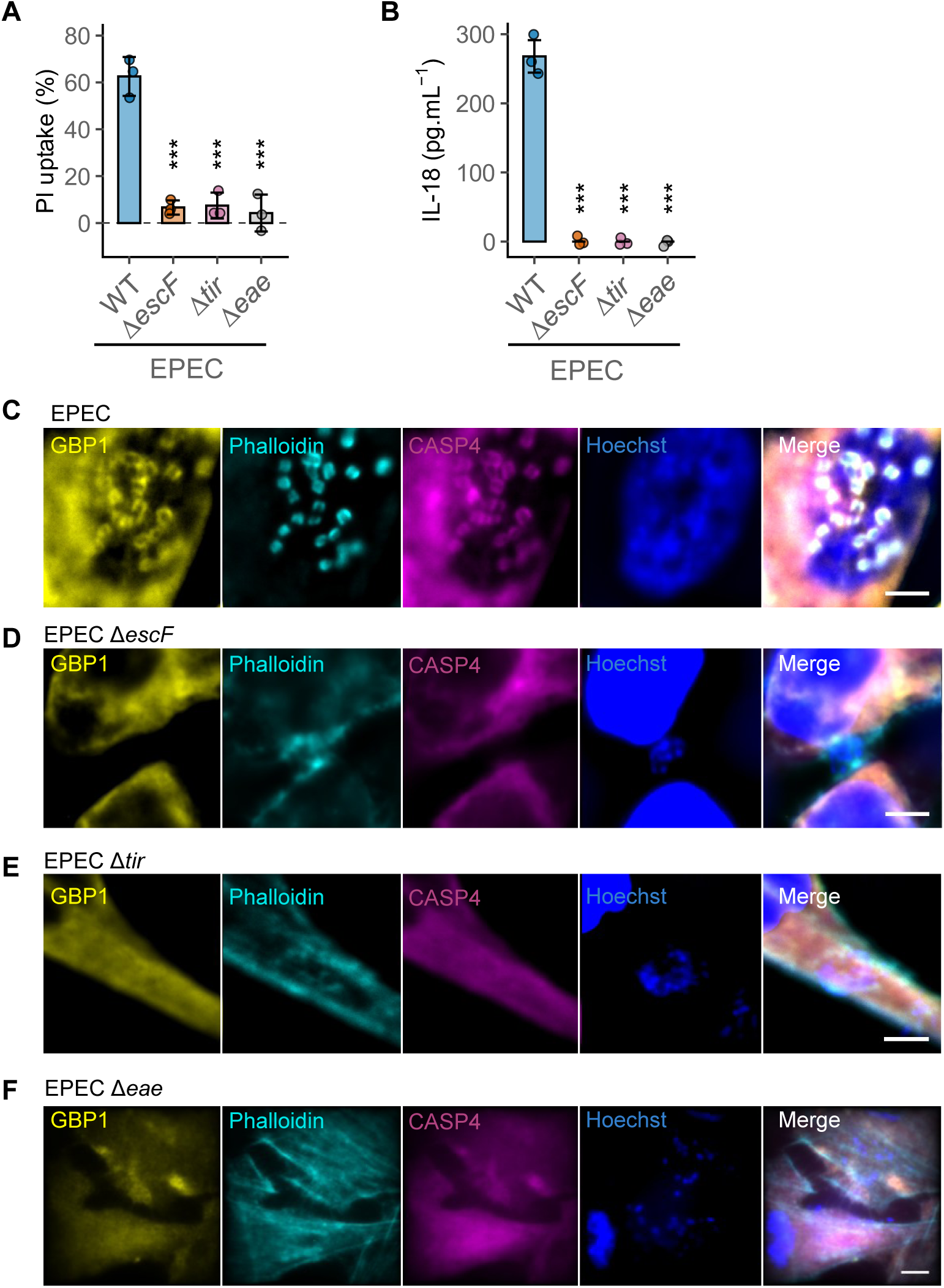
GBP1 recruitment and pyroptosis requires Tir-Intimin signalling leading to actin polymerisation. (A) – (B) Percentage pyroptotic cell death as measured by propidium iodide dye uptake assays (A) or IL-18 quantified by ELISA (B) from IFNγ-primed HeLa cells infected with the wild-type (WT) EPEC or the indicated mutants for 6 h. Mean ± SD error bars with symbols representing data from n = 3 independent experiments are shown. In (A) *\*\*\* P = 0.0002* and in (B) *** *P = 1.4e-07* are two-tailed *P* values for comparisons between the WT and indicated mutants from mixed effects ANOVAs. (C) – (F) Representative immunofluorescence images of IFNγ-primed HeLa cells expressing YFP-Caspase4^C285S^ infected with WT EPEC or the indicated mutants for 2 h. Cells were stained with anti-GBP1 antibodies, phalloidin (to stain actin) and Hoechst (DNA dye; blue). Data from n = 3 independent experiments. Scale bar, 5 μm.

### GBP1 mutants with LPS-binding defects can traffic to EPEC pedestals

We next focussed on discovering the biochemical features of GBP1 that govern its trafficking to bacterial pedestals. The GTP-binding deficient K51A mutant and farnesylation-deficient C589A, which are required for association with LPS and formation of polymeric coats (<u>Kutsch & Coers, 2021</u>; <u>Rivera-Cuevas *et al*., 2023</u>; <u>Tretina *et al*., 2019</u>) and could not restore pyroptosis defects in *GBP1^-/-^* cells (***Figure 1C***), also failed to traffic to EPEC pedestals (***Figure EV5A, 4A***). This suggests that GBP1 localisation at actin-rich pedestals is required for pyroptosis. As A/E pathogens remain extracellular, we asked whether residues in GBP1 known to contribute to LPS binding were required for its trafficking to pedestals. Farnesylated GBP1 transitions from a “closed” head-to-head dimer to crossover dimers with an “open” extended conformation on purified LPS membranes and natively on Gram-negative bacteria (***Figure EV1C***) (<u>Kuhm *et al*., 2025</u>; <u>Weismehl *et al*., 2024</u>; <u>Zhu *et al*., 2024</u>). Mutation of residues in the linker joining the large GTPase domain (LG) and middle domain (MD) that enable parallel crossover dimers and those within the LG domain required for GTPase activity and polymerisation severely attenuate GBP1 LPS attachment and formation of bacterial coats (<u>Kuhm *et al*., 2025</u>; <u>Weismehl *et al*., 2024</u>; <u>Zhu *et al*., 2024</u>). We therefore expressed two GBP1 variants: GBP1^Δlinker^ (^308^DLP^310^⃗AAA mutations in the LG-MD linker; (<u>Kuhm *et al*., 2025</u>); ***Figure EV1C, S5B***) and GBP1^M139D^ (within the LG domain that lacks GTPase and oligomerisation activity; (<u>Weismehl *et al*., 2024</u>); ***Figure EV1C, S5B***) and assessed their localisation to EPEC pedestals. Immunofluorescence microscopy revealed that WT GBP1 and both mutants were recruited to similarly to EPEC pedestals (***Figure 4A-B***). However, unlike WT GBP1, and consistent with their inability to bind to LPS, both variants failed to complement the pyroptosis defects of *GBP1^-/-^* cells upon infection with EPEC (***Figure 4C***). Taken together, we conclude that GTP-binding and prenylation are required for GBP1 targeting to actin-rich pedestals, but its recruitment to these sites proceeds independently of its ability to detect LPS.

**Figure 4.**
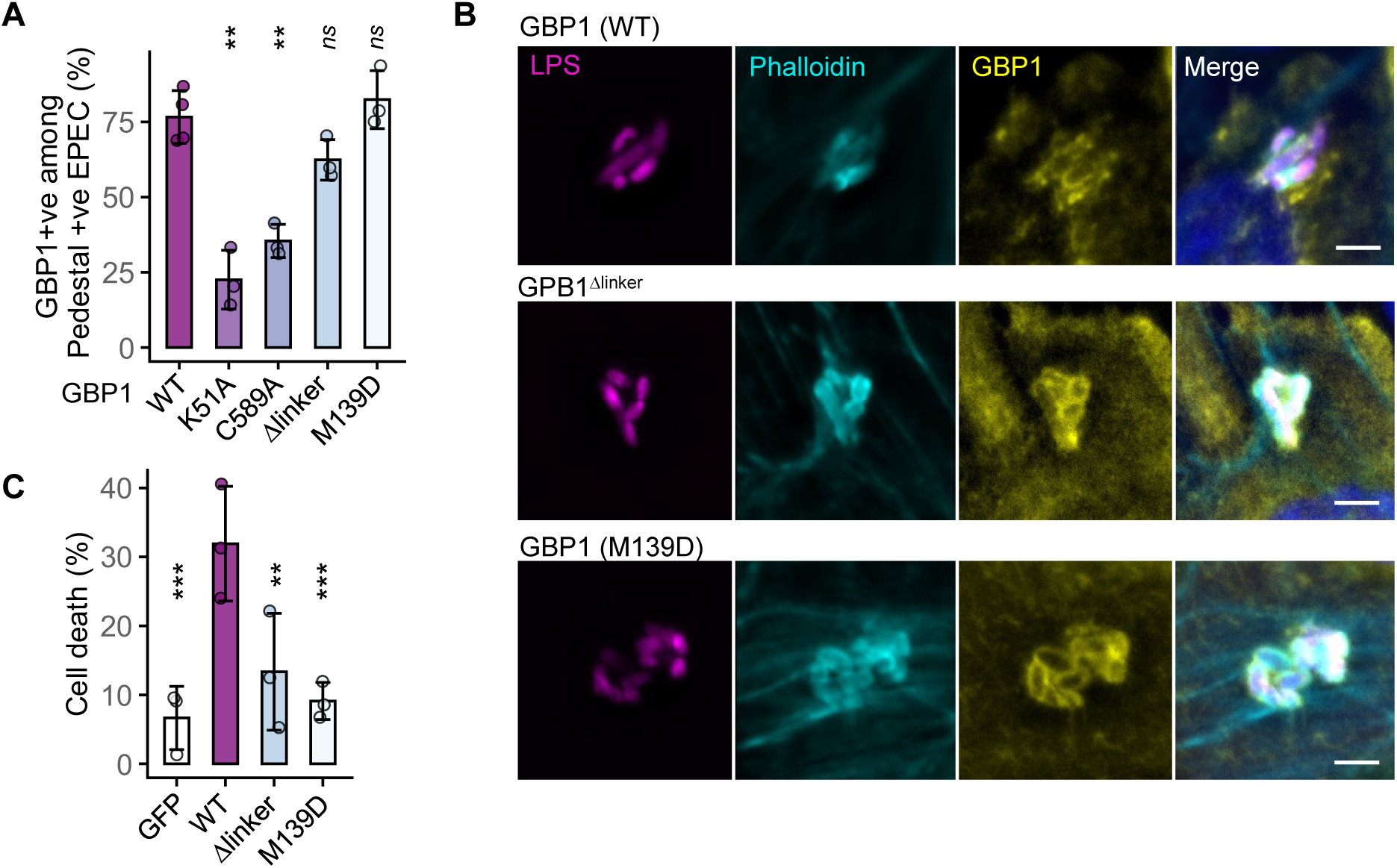
GBP1 mutants defective for LPS binding are recruited to EPEC actin-rich pedestals. (A) Quantification of EPEC pedestals in IFNγ- and doxycycline-treated HeLa *GBP1^-/-^* cells with Tet-inducible expression of the indicated wild-type (WT) or mutant GBP1 variants that stained positive for GBP1 at 3 h post-infection. Mean ± SD error bars with symbols representing individual fields of view from n = 3 independent experiments are shown. *\*\* P < 0.01, ns –* not significant *(P > 0.05)* are two-tailed *P* values for comparisons between WT GBP1 and other groups from mixed effects ANOVA. Exact *P* values as follows – K51A: 0.0008; C589A: 0.0034; Δlinker: 0.1533; M139D: 0.316. (B) Representative images from IFNγ- and doxycycline-treated HeLa *GBP1^-/-^* cells with Tet-inducible expression of WT GBP1 or the indicated variants. Cells were stained with anti-LPS (to stain bacteria) and anti-GBP1 antibodies, phalloidin (to stain actin) and Hoechst (DNA dye; blue). Data are presentative of n = 3 independent experiments. Scale bar, 4 μm. (C) Percentage pyroptotic cell death as measured by propidium iodide dye uptake assays from IFNγ- and doxycycline-treated *GBP1*^-/-^ HeLa cells expressing Tet-inducible GFP (as negative control) or the WT or indicated mutants of GBP1. Mean ± SD error bars with symbols representing data from n = 3 independent experiments are shown. *\*\* P < 0.01, *** P < 0.001* are two-tailed *P* values for comparisons between GBP1 and other groups from mixed effects ANOVA. Exact *P* values as follows – GFP: 0.0005; Δlinker: 0.0011; M139D: 0.0005.

### GBP1 is recruited to ‘sterile’ actin polymerisation sites independently of LPS

To further verify LPS-independent trafficking of GBP1 to bacterial actin-rich pedestals, we devised a novel, bacteria- and LPS-free assay that involves Tir-dependent actin polymerisation. We note that the signal transduction by Tir^EPEC^ is analogous to eukaryotic immunoglobulin (Ig) Fragment crystallisable (Fc) receptors (FcR) because the crosslinking of the extracellular ligand-binding regions of both receptors triggers intracellular signalling via phospho-tyrosine motifs (<u>Pincetic *et al*, 2014</u>). We therefore engineered a chimeric receptor composed of the signal peptide and the extracellular IgG-binding domain of the human Fcγ receptor IIA and the intracellular C-terminal region of Tir^EPEC^, which is sufficient for downstream signalling to N-WASP-ARP2/3 and actin polymerisation (<u>Frankel *et al*., 1998</u>) (i.e., FcγR-Tir fusion protein; ***Figure 5A***). We reasoned that crosslinking of chimeric FcγR-Tir with human serum IgG (hIgG)-coated polystyrene beads would lead to clustering of the Tir intracellular domain leading to actin polymerisation (***Figure 5A***).

**Figure 5.**
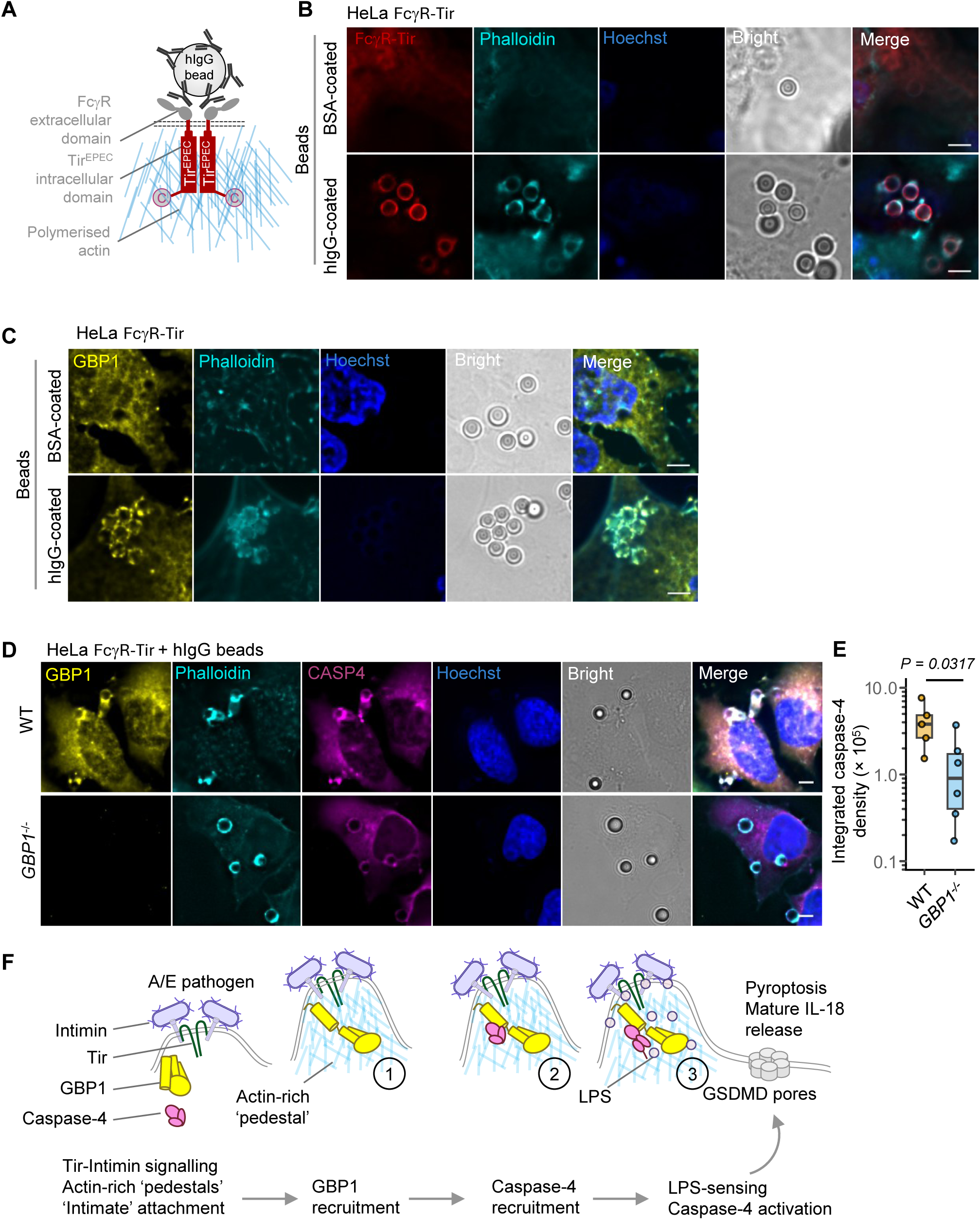
GBP1 is recruited to ‘sterile’ sites of actin polymerisation independently of LPS. (A) A schematic showing the reconstitution of Tir-driven actin polymerisation with an FcγR-Tir chimeric fusion protein containing the extracellular ligand-binding (IgG-binding) domain of FcγRIIA and the intracellular C-terminal region of Tir^EPEC^ that is sufficient for actin-polymerisation. Clustering of FcγR-Tir with sterile polystyrene beads (3.54 μm diameter) coated with human IgG (hIgG) results in actin-rich structures at sites of bead attachment. The FcγR-Tir chimaera with a C-terminal mCherry2 tag used for imaging in (B) or a non-fluorescent MYC tag in (C)-(D) behaved similarly (also see ***Figure EV5***). (B) Representative images from HeLa cells expressing FcγR-Tir (with C-terminal mCherry2 tag) treated for 3 h with sterile polystyrene beads coated with bovine serum albumin (BSA) or hIgG as labelled. Cells were stained with phalloidin (to stain actin) and Hoechst (DNA dye; blue) and mCherry2 fluorescence showed FcγR-Tir localisation. Data are presentative of n = 3 independent experiments. Scale bar, 5 μm. (C) Representative images from HeLa cells expressing FcγR-Tir treated for 3 h with sterile polystyrene beads coated with BSA or hIgG. Cells were stained with anti-GBP1 antibody, phalloidin (to stain actin) and Hoechst (DNA dye; blue). Data are presentative of n = 3 independent experiments. Scale bar, 5 μm. (D) Representative images from wild-type (WT) or *GBP1*^-/-^ HeLa cells expressing FcγR-Tir and YFP-Caspase-4^C258S^ treated for 3 h with hIgG-coated sterile polystyrene beads. Cells were stained with anti-GBP1 antibody, phalloidin (to stain actin) and Hoechst (DNA dye; blue). Data are presentative of n = 4 independent experiments. Scale bar, 5 μm. (E) Local caspase-4 intensity around the sites of actin-rich structures in WT or *GBP1*^-/-^ HeLa cells expressing the FcγTir fusion protein. Quantification of 5 (WT) or 6 (*GBP1*^-/-^) independent fields of view across at least 3 biological replicates was measured using ImageJ. Box plots, median and interquartile range (IQR) are shown, with boxes indicating IQR, whiskers indicating 1.5x IQR and the line indicating the median. Two-tailed *P* value for the indicated comparison from an independent sample Student’s *t* test is indicated. (F) Schematic summary of EPEC/EHEC-induced pyroptosis and IL-18 production in three-steps (indicated in numbers inside circles) and briefly outlined in the text below. The initial attachment and microcolony formation (as in the case of EPEC) leads to the translocation of Tir and its clustering by Intimin expressed on the bacterial surface. In step 1, GBP1 is recruited by pathogen-induced actin polymerisation, in step 2 caspase-4 is recruited in a GBP1-dependent manner, and in Step 3, caspase-4 is activated by LPS at sites of focal bacterial attachment, leading to pyroptosis via gasdermin D (GSDMD) cleavage and IL-18 maturation.

HeLa cells expressing FcγR-Tir challenged with hIgG-coated beads showed clustering of the FcγR-Tir chimaera and dense actin polymerisation around bead-attachment sites (***Figure 5B***). No FcγR-Tir clustering or actin polymerisation was observed with BSA-coated beads used as a negative control (***Figure 5B***). As an additional control, we developed FcγR-Tir^mut^ (with point mutations in the tyrosine residues in the intracellular domain of Tir that abolish actin polymerisation), which also failed to trigger actin polymerisation upon clustering with hIgG-coated beads (***Figure EV5C***). Similarly, treatment with cytochalasin D, an actin polymerisation inhibitor, also blocked the formation of actin structures induced by hIgG-beads (***Figure EV5D***). Together, this confirmed that signalling by FcγR-Tir is induced only upon clustering with antibody ligands leading to actin polymerisation through intracellular amino acid motifs known to be important for signalling in native Tir^EPEC^ when clustered by intimin.

We next assessed endogenous GBP1 localisation in this experimental setting that produces ‘sterile’ actin-rich structures independently of bacteria or LPS. GBP1 was prominently recruited to actin-rich structures produced in FcγR-Tir cells treated with hIgG-beads but not BSA-beads (***Figure 5C***). GBP1 recruitment was blocked when actin polymerisation was abrogated in cells expressing FcγR-Tir^mut^ or upon cytochalasin D treatment (***Figure EV5C-D***). These experiments provided genetic and pharmacological evidence that actin polymerisation that mimics A/E pathogens is sufficient to elicit GBP1 mobilisation.

Similar to native bacterial infection, caspase-4 localised to sterile actin-rich structures produced in FcγR-Tir cells (***Figure 5D***), indicating that caspase-4 can also traffic to pedestals independently of LPS. Despite similar levels of actin-rich structures in *GBP1*^-/-^ FcγR-Tir cells treated with hIgG-beads, caspase-4 localisation at these structures was markedly reduced in *GBP1*^-/-^ cells; this was verified by quantification of caspase-4 intensity within the actin-rich structures around bead-attachment sites (***Figure 5D-E***). This implies that GBP1 is recruited to bacterial attachment sites independently of LPS and facilitates caspase-4 recruitment. As FcγR-Tir cells treated with BSA or hIgG beads for 6 h did not undergo appreciable levels of pyroptosis (***Figure EV5E***), we inferred that GBP1-caspase-4 recruitment to actin-rich structures is not sufficient for triggering cell death, presumably due to the absence of LPS to stimulate caspase-4 proteolytic activity in this setting. We therefore conclude that GBP1-dependent and LPS-independent caspase-4 recruitment to actin-rich pedestals of A/E pathogens permits the detection of bacterial LPS at sites of focal bacterial attachment (***Figure 5F***).

## Discussion

Here we discovered that GBP1 responds to A/E bacteria-induced actin polymerisation and its effectiveness in surveillance of extracellular pathogen niches. GBP1 was recruited to ‘sterile’ actin polymerisation sites induced using our novel FcγR-Tir chimeric receptor, revealing that its mobilisation to bacterial attachment sites is LPS-independent. Mutation of residues in GBP1 that enable LPS-sensing did not affect its trafficking to EPEC, suggesting that its ETI activity is separate from its PRR function. GBP1 also mediated caspase-4 recruitment in an LPS-independent manner. During A/E pathogen infection, GBP1-dependent caspase-4 recruitment leads to pyroptotic host-cell death and IL-18 release (***Figure 5F***). These findings add a new dimension to the antimicrobial actions of GBP1 which was previously characterised as a cytosolic PRR for LPS.

How LPS activates caspase-4/11 is of much interest given the highly inflammatory outcomes associated with pyroptosis (<u>Hagar *et al*, 2013</u>; <u>Kayagaki *et al*, 2013</u>). EPEC/EHEC and other Gram-negative bacteria shed LPS-containing outer membrane vesicles (OMVs) that can be internalised by host cells (<u>Croxen *et al*, 2013</u>; <u>Kulp & Kuehn, 2010</u>; <u>Vanaja *et al*, 2016</u>). We previously showed that LPS enters IFNγ-primed human epithelial cells infected with EPEC in a manner that depends on Ca^2+^ influx through the TRPV2 ion channels, which is induced by Tir clustering by Intimin (<u>Zhong *et al*., 2022</u>; <u>Zhong *et al*., 2020</u>). Here we found that actin polymerisation by Tir recruits GBP1 to pedestals, which is followed by caspase-4 for LPS-sensing at bacterial attachment sites. We and others have previously showed that GBP1 and caspase-4 form a complex in response to bacterial infection or LPS transfection (<u>Fisch *et al*., 2019</u>; <u>Kutsch *et al*., 2020</u>; <u>Santos *et al*., 2020</u>; <u>Wandel *et al*., 2020</u>), therefore they may also interact during EPEC/EHEC infection leading to their localization to bacterial pedestals. Further, murine caspase-4/11 is proposed to localise to the leading edge in leukocytes; therefore, the diffuse caspase-4 staining at EPEC/EHEC pedestals in *GBP1*^-/-^ cells may point to its ability to traffic to the plasma membrane in homeostasis (<u>Li *et al*, 2007</u>). We propose that the pyroptosis induced by extracellular A/E pathogens proceeds via three steps: (i) trafficking of GBP1 to actin-rich pedestals to ‘prime’ the response, (ii) recruitment of caspase-4 for LPS-sensing, and (iii) stimulation of caspase-4 activity leading to pyroptosis and cytokine maturation (***Figure 5F***).

GBP1 mobilisation to EPEC/EHEC pedestals promoted pyroptosis, thus establishing its role in ETI against these extracellular pathogens. The loss of GBP1 prenylation (GBP1^C589A^), GTP-binding (GBP1^K51A^), GTPase activity (GBP1^M139D^) and compromised crossover dimer formation (GBP1^Δlinker^) abolished its pyroptotic functions during EPEC infection, indicating that GBP1 membrane recruitment, GTPase activity and higher-order oligomerisation are essential for A/E pathogen-induced pyroptosis. We reason that GBP1 may operate like the Ras-family of small GTPases which also contain C-terminal prenylation signals and polybasic motifs that enable GTP-activity-dependent signal transduction (<u>Jung & Bachmann, 2023</u>; <u>Williams, 2003</u>). GBP1 recruitment to bacterial attachment sites was strictly dependent on actin polymerisation but independent of the downstream signalling mechanism used by EPEC and EHEC (tyrosine phosphorylation by Tir^EPEC^ and TccP by Tir^EHEC^; ***Figure EV4A***). The signal(s) that induce GBP1 trafficking to Tir-dependent actin-rich structures are not known. We ruled out the involvement of IRTKS or IRSp53 in recruiting GBP1, because both proteins are recruited to Tir during infection by the EHEC Δ*tccP* strain which did not recruit GBP1 (<u>Naydenov *et al*, 2024</u>). We speculate that GBP1 recruitment may involve direct interactions with actin, which is a known binding-partner of GBP1 (<u>Ostler *et al*, 2014</u>), sensing of altered membrane curvature generated by I-BAR (Inverse – Bin1, amphiphysin, Rvs167) domain-containing proteins recruited by Tir (<u>Naydenov *et al*., 2024</u>; <u>Zhao *et al*, 2011</u>), phosphoinositides generated during Tir-signalling (<u>Sason *et al*, 2009</u>), or sugars, which have been proposed to recruit human GBP1 and mGbp2 to damaged vacuoles (<u>Feeley *et al*, 2017</u>).

We also uncovered biochemical differences between the ETI role of GBP1 and previously reported features that support its LPS PRR activities. For instance, here we observed GBP1 trafficking to actin-rich pedestals of pathogenic *E. coli* strains expressing distinct O-antigens. However, GBP1 trafficking to a rough LPS mutant of *S. flexneri* (that lack an O-antigen) is markedly reduced (<u>Piro *et al*., 2017</u>), suggesting that the O-antigen only influences the PRR activity of GBP1. More importantly, we established that GBP1 trafficking to A/E pathogen attachment sites was completely LPS-independent as demonstrated by its recruitment to actin-rich structures generated using the FcγR-Tir chimeric receptor. Mutational analyses further indicated that the PRR activity of GBP1 (LPS interaction) and ETI activity (trafficking to pedestals) are separable. The GBP1^K51A^ mutant cannot bind GTP, dimerise or form higher-order multimers, but the GBP1^M139D^ variant can bind GTP but cannot hydrolyse it and fails to dimerise or form polymeric coats. The GBP1^Δlinker^ mutant is compromised in crossover dimer formation in the extended LPS membranes/bacteria-bound conformation. Taken together, we conclude that GTP-binding of farnesylated GBP1 promotes its trafficking to A/E pathogen pedestals, and dimerization/polymerisation may be dispensable.

Whether GBP1-dependent caspase-4 recruitment to Tir-driven structures is fundamentally distinct from its recruitment to the surface of *Salmonella/Shigella* awaits further investigation. GBP1 can detect LPS-free intact pathogen-containing vacuoles (e.g., *Toxoplasma*), which could serve as a model to dissect whether GBP1 can recruit caspase-4 to other intracellular pathogen niches. However, stimulation of caspase-4 activity would still require LPS; therefore, any non-canonical roles for caspase-4 in such scenarios also remain to be identified (<u>Abu Khweek & Amer, 2020</u>).

In contrast to the role of GBP2-5 in responding to cytosolic bacteria, GBP1 appears to play a dominant role against extracellular A/E pathogens. Although GBP2 was occasionally found in the vicinity of EPEC pedestals, *GBP2* silencing did not alter pyroptosis; therefore, future work should investigate pyroptosis-independent cell-intrinsic roles of human GBP2 against A/E pathogens. Surprisingly, EPEC-induced pyroptosis could be not be sustained by GBP2 expression in *GBP1*^-/-^cells, even though GBP2 can redundantly serve as an LPS PRR during *S. flexneri* infection of *GBP1*-deficient cells by aggregating LPS OMVs shed by the bacterium directly in the cytosol (<u>Dickinson *et al*., 2023</u>). We posit that relatively low levels of LPS are internalised during infection by extracellular EPEC/EHEC, which requires GBP1-dependent actions due to its high affinity for LPS (dissociation constant ∼60 nM (<u>Santos *et al*., 2020</u>)). Consistent with this, 100-times greater LPS concentration is required for its aggregation *in vitro* by GBP2 as compared to GBP1, pointing to the much weaker affinity of GBP2 for LPS (<u>Dickinson *et al*., 2023</u>).

It is likely that the high affinity of GBP1 for LPS is also the reason that its targeting to polymerised actin is context dependent. For instance, GBP1/mGbp2 do not target actin ‘comets’ generated by cytosolic *Shigella* and *Burkholderia* (<u>Dilucca *et al*, 2021</u>; <u>Piro *et al*., 2017</u>; <u>Place *et al*, 2020</u>; <u>Wandel *et al*, 2017</u>). In these scenarios, the abundant cytosolic LPS may be the dominant signal leading to the formation of coats directly on bacteria. Alternatively, because GBP1 is a farnesylated GTPase, it may preferentially target pathogen-induced cortical polymeric actin structures as opposed to cytosolic actin comets. Similarly, GBP1 does not target the cytosolic actin comets of *Listeria monocytogenes*; in contrast, GBP1 targets membrane remnants following the escape of *Listeria* from its vacuole into the cytosol, pointing to exposed sugars from the vacuolar lumen as alternative signals that can attract GBP1 (<u>Buijze *et al*, 2023</u>; <u>Kim *et al*., 2011</u>; <u>Santos *et al*., 2020</u>). The mechanisms of actin-based motility vary markedly among bacterial pathogens, which may underlie the lack of GBP1 targeting to bacteria. For instance, A/E pathogens (Tir), *Shigella* (IcsA)*, Burkholderia* (BimA)*, Rickettsia* (RickA, Sca2), *Listeria* (ActA) differentially coopt actin nucleation promoting factors such as N-WASP, ARP2/3, ENA/VASP and others (<u>Choe & Welch, 2016</u>). Whether GBP1 is recruited to other pathogens that manipulate cortical actin, such as vaccinia virus or *Cryptosporidium*, will be worth investigating in the future. As GBP1 recruited caspase-4 to FcγR-Tir-induced actin-rich structures, comparative studies will also help dissect niches to which caspase-4 is recruited by GBP1, for example to pathogen-containing vacuoles, and clarify whether A/E pathogen pedestals present a distinct of mode of caspase-4 engagement by GBP1.

During CR infection of mice, caspase-4–dependent IL-18 production contributes to neutrophil influx in the gut, which can be antagonised by the CR caspase-11/4 inhibitory effector NleF early during infection (up to day 4 of CR infection) (<u>Pallett *et al*, 2017</u>). The production of IFNγ by the host protects against CR and may help overcome the suppressive effect of NleF by upregulating mGbp2 expression (<u>Mullineaux-Sanders *et al*, 2021</u>; <u>Simmons *et al*, 2002</u>), which could be tested in *Gbp2-*deficient mice in the future. EPEC and EHEC have convergently evolved mechanisms for actin polymerisation, pointing to its important role in pathogenesis. Additionally, unlike the prototypical EPEC 2348/69, many typical and atypical strains of EPEC express TccP and/or the highly related TccP2 and deploy both actin polymerisation pathways, hinting at strong selection for actin polymerisation among A/E pathogens (<u>Martins *et al*, 2017</u>). The remodelling of the actin cytoskeleton disrupts tight junctions and barrier function in the gut and promotes pathogenesis (<u>Collins *et al*., 2014</u>). It is tempting to speculate that GBP1 has therefore evolved a mechanism to detect pathogen-induced actin polymerisation, which is the final result of EPEC/EHEC attachment, and trigger pyroptosis to clear away cells with adherent pathogens. This may provide a failsafe against EPEC/EHEC as their flagellins and T3SS systems evade detection by the NLRC4 inflammasomes which can readily detect similar molecules of *Salmonella* and *Shigella* (<u>Sanchez-Garrido *et al*., 2020</u>; <u>Shenoy *et al*., 2018</u>).

In summary, here we uncovered a novel interface between actin cytoskeleton manipulation and innate immunity via GBP1, i.e., LPS-independent GBP1-trafficking to sites of actin polymerisation induced by extracellular bacteria. The GBPs and caspase-4-like enzymes are conserved across vertebrate species, including domestic animals and pets that can be infected by A/E pathogens (<u>Corte-Real *et al*, 2022</u>; <u>Devant *et al*, 2021</u>; <u>Digby *et al*, 2021</u>; <u>Schelle *et al*, 2023</u>). However, LPS-sensing is not universally conserved among caspase-4-related enzymes (<u>Digby *et al*., 2021</u>). Similarly, whether GBP1-like proteins share LPS-sensing responses should also be assessed given the lineage-specific expansion and loss of the *GBP* gene cluster in vertebrates. The cytoskeleton is crucial for homeostasis and host defence (<u>Mostowy & Shenoy, 2015</u>) and a plethora of pathogens or their secreted toxins that hijack the actin cytoskeleton (<u>Aktories *et al*, 2011</u>; <u>Stevens *et al*, 2006</u>). The role of GBP1 in ETI against cytoskeleton manipulation and whether it responds to actin polymerisation in homeostasis are additional question to be addressed in the future. Our discovery that GBP1 can respond to extracellular pathogens markedly broadens its scope as a sentinel of microbial infection.

## Methods

### Reagents, cell lines and bacterial strains

Key reagents, antibodies, cell lines and bacterial strains are listed in ***Reagents and Tools Table***.

### Cell culture

Cell lines used in this study are listed in ***Reagents and Tools Table***. Cells were maintained in a humidified incubator at 37 °C with 5 % CO_2_ and verified to be free from mycoplasma. Cells were sourced from ATCC and tested negative for mycoplasma. HeLa and HEK293E cells were cultured in complete Dulbecco’s Modified Eagle’s Medium (DMEM) containing 4500 mg.L^-1^ glucose, 100 units (U).mL^-1^ penicillin, 100 µg.mL^-1^ streptomycin, 1 mM sodium pyruvate and 10 % (v/v) heat-inactivated foetal calf serum (HI-FCS). CL40 cells were grown in DMEM/F12 Ham (with 100 U.mL^-1^ penicillin, 100 µg.mL^-1^ streptomycin, 1 mM sodium pyruvate and 20 % (v/v) HI-FCS). Where required, cultures were supplemented with puromycin (2 µg.mL^-1^), blasticidin S (10 µg.mL^-1^) or zeocin (200 µg.mL^-1^). Prior to experiments, cells were primed with 10 ng.mL^-1^ IFNγ for 16 h in medium without penicillin, streptomycin and phenol red, and where required, protein expression was induced by adding 200 ng.mL^-1^ doxycycline along with IFNγ.

### Growth of bacterial strains

All bacterial strains used in this study are listed in ***Reagents and Tools Table***. Bacteria were routinely grown overnight in lysogeny broth (LB) at 37 °C with shaking (180 rpm). Appropriate antibiotics were added when necessary: ampicillin (100 µg.mL^-1^); kanamycin (30 µg.mL^-1^), chloramphenicol (20 µg.mL^-1^). Infections were carried out using DMEM-primed EPEC or EHEC, resulting in increased expression of LEE and non-LEE encoded virulence factors as described before (<u>Goddard *et al*., 2019</u>; <u>Zhong *et al*., 2022</u>; <u>Zhong *et al*., 2020</u>). For EPEC-priming before infections, overnight cultures grown in LB were diluted 1:50 into pre-equilibrated DMEM-low glucose (1000 mg.mL^-1^ glucose; Sigma D5546) supplemented with antibiotics as required, and incubated for 3 h in a humidified incubator at 37 °C with 5 % CO2. For EHEC-priming, cultures were grown in LB for 6 h and then in DMEM-low glucose for 18 h in a humidified incubator at 37 °C with 5 % CO_2_.

### Bacterial infections

DMEM-primed bacterial cultures were suspended in fresh DMEM at desired multiplicity of infection (MOI) following quantification by measuring optical density at 600 nm. Epithelial cell lines pre-treated with IFNγ were infected with bacteria at an MOI of 5 for microscopy or 30 for cell death assays (verified by counting CFU), unless otherwise stated. Infections were synchronised by centrifugation at 750 x*g* for 10 min. Typical EPEC form microcolonies during infection whereas EHEC does not and therefore takes longer to attach tightly to cells and translocate effectors, including Tir. EPEC infections were carried out for 2 h and EHEC infections for 5 h before the addition of gentamicin (200 µg.mL^-1^) to kill extracellular bacteria. For pyroptosis assays described below, EPEC infections were stopped at 6 h post-infection (hpi) in HeLa and CL40 cells or 12 h for HT29 cells and EHEC infections at 8 hpi as indicated in figure legends. For microscopy, EPEC infection was stopped at 2 hpi and EHEC infection at 5 hpi, followed by fixation and processing as described below.

### Infection of mice with *Citrobacter rodentium* (CR)

Mouse experiments were conducted with five mice per group. Pathogen-free female 18-20 g C57BL/6 mice were purchased from Charles River Laboratories. All mice were housed in pathogen-free conditions, at 20-22°C, 30-40 % humidity on 12 h of light/dark cycle in high-efficiency particulate air (HEPA)-filtered cages with sterile corn cob bedding, nesting material, and enhancements (chewing toy, and opaque and transparent cylinders), and were fed with RM1 (E) rodent diet (SDS diet) and water ad libitum.

CR strain ICC169 (Nalidixic acid (Nal) resistant) (<u>Schauer & Falkow, 1993</u>) was grown overnight in LB supplemented with 50 μg.ml^-1^ nalidixic acid at 37 °C at 180 rpm, were centrifuged at 3000 x*g* for 10 min, and resuspended in sterile 1X phosphate buffered saline (PBS). Mice weight were recorded (d0) and they were infected with approximately 3 x 10^9^ CFUs in 200 μl sterile PBS by oral gavage as previously described (<u>Crepin *et al*, 2016</u>). For mock infection (uninfected mice), mice received 200 μl sterile PBS. The inoculum CFUs were confirmed by CFU quantification as previously described (<u>Crepin *et al*., 2016</u>). Infections were followed by plating mouse stools at 2-3 days post infection (dpi) onwards on LB agar (15 % v/v) + Nal plates as previously described (<u>Crepin *et al*., 2016</u>).

### Immunohistostaining of colonic sections

Mouse colons were harvested at 8 dpi, fixed in 4 % paraformaldehyde (PFA) for 2.5 h, and immersed in 70 % ethanol. The fixed tissues were embedded in paraffin and sectioned at 5 μm. For immunofluorescence, sections were dewaxed by immersion in Histoclear solution twice for 10 min, followed by immersion in 100 % ethanol for 10 mins, twice, then 95 % ethanol for 3 min twice, 80 % ethanol for 3 min, and 1X PBS-0.1 % Tween 20, 0.1 % saponin (PBS-TS), for 3 min twice. The sections were heated for 30 min in demasking solution (0.3 % trisodium citrate, 0.05 % Tween-20 in distilled H_2_O). The slides were washed in PBS-TS, followed by blocking in PBS-TS supplemented with 10 % normal donkey serum (NDS) for 20 min. The slides were incubated overnight at 4 °C with mouse anti-CR polyclonal antibody (1:50; (<u>Mullineaux-Sanders *et al*., 2021</u>)), rabbit anti-GBP2 polyclonal antibody (1:100). The following day the slides were washed twice for 10 mins in PBS-TS, and incubated with the appropriate secondary antibody (1:100) and DAPI (1:1000) to stain DNA (***Reagents and Tools Table***). The slides were washed and mounted with ProLong Gold antifade mountant.

### Cell death assays and ELISAs

For propidium iodide (PI)-uptake assays, cells were seeded in 96-well black-wall clear-bottom plate at a seeding density of 2 x 10^4^ cells/well in 100 µL complete DMEM lacking antibiotics and phenol red (to reduce background fluorescence) (<u>Goddard *et al*., 2019</u>). Cells were primed for 16 h with 10 ng.mL^-1^ IFNγ or as specified in the figure legend. The medium was supplemented with 5 µg.mL^-1^ PI prior to infection and values in uninfected cells was used as baseline, which was subtracted from all values. Proportional uptake of PI was calculated as a percentage relative to the baseline-corrected values of uninfected cells given 0.05 % (v/v) Triton X-100. Infections were carried out at 37 °C with 5 % CO_2_ using a FLUOStar Omega microplate reader (BMG Labtech). Fluorescence measurements were taken using 540/10 excitation and 620/10 emission filters.

For ELISAs, after infection, cells were centrifuged at 750 x*g* for 10 mins to collect supernatants. IL-18 was quantified using the Human Total IL-18 DuoSet ELISA kit (R&D Systems) according to the manufacturer protocol, using 96-well high-binding ELISA plates (Greiner). Samples were measured using a FLUOStar Omega microplate reader using absorbance at 450 nm, and corrected by subtracting absorbance at 540 nm.

### Generation of *GBP1-*knockout cells

HeLa cells seeded in 24-well plates at 1 x 10^5^ cells/well were co-transfected with 125 ng each of two plasmids encoding both constitutively expressed *Sp*Cas9 and *GBP1*-specific gRNA using TransIT-X2 Dynamic Delivery System (Mirus) as per the manufacturer guidelines, with transfection complexes prepared in Opti-MEM (Gibco). The gRNA sequences were as follows: GBP1 sgRNA1: CTCATAAGCTGGTACCACTC; GBP1 sgRNA2: TACATACAGCCAGGATGCAA. Together, these gRNAs aimed to remove the entire coding sequence of *GBP1*. Puromycin was added to the wells at a final concentration of 2 µg.mL^-1^ at 24 h post-transfection to select for transfected cells, and selection was continued for 72 h. Surviving cells were then trypsinised using 0.25 % trypsin-EDTA (Sigma), and limited dilutions were performed to a final concentration of 1 cell.mL^-1^ in complete DMEM to obtain isolated clonal islands upon plating in a 10 cm tissue-culture treated dish (Greiner). To confirm *GBP1* deficiency on the genetic level, the target locus was subject to Sanger sequencing, and GBP1 expression was assessed using western blotting.

### Stable silencing with miRNA30E

For stable knock-downs of target genes, 22-base oligonucleotides were cloned into the pMX-CMV-eYFP-miR30E backbone as described previously for caspase-4 (<u>Eldridge *et al*, 2017</u>; <u>Fellmann *et al*, 2013</u>). Briefly, 22 base oligonucleotides specific to the gene of interest were used in the enhanced miR30E version. The sequences are listed in ***Reagents and Tools Table***. Using Sequence and Ligation Independent Cloning (SLIC) (<u>Jeong *et al*, 2012</u>), these 22mer sequences were cloned into the retroviral pMXCMV-YFP-MCS plasmid at the XhoI and EcoRI sites. pMXCMV-miR30E plasmids were then transduced into the recipient cell line as described below and expression of the gene of interest analysed via western blotting relative to the non-targeting control plasmid (***Reagents and Tools Table***). If required, eYFP-positive cells were selected via FACS using an ARIA III (BD Bioscience).

### Molecular biology methods for generating mutants and domain swaps

Throughout this study, SLIC was used for routine cloning, and Phusion polymerase or KOD hot-start polymerase were used for PCR for their high fidelity (<u>Jeong *et al*., 2012</u>). Oligonucleotides used are listed in ***Reagents and Tools Table***, and all plasmid constructs were confirmed using Sanger sequencing (Azenta). Expression of GBP1 was carried out using derivatives of the doxycycline-inducible template vector pTetDF-mCh or similar in conjunction with the control vector pTet-rtTA-tTS (<u>Fisch *et al*., 2019</u>). Human FcγRIIA (aa 1-245) extracellular Fc-binding region and the Tir^EPEC^ (aa 387-550, either WT sequence or with Y454A and Y474A mutations to generate an actin polymerisation-defective FcγR-Tir^mut^) (<u>Campellone & Leong, 2005</u>) and either mCherry2 or 2x MYC tag (GSEQKLISEEDLEQKLISEEDL) were fused by PCR and cloned into the retroviral packaging plasmid pMXCMV.

### Packaging and transduction of retro- and lenti-viral vectors

Virus-like particles were packaged and produced in HEK293E cells as described previously (<u>Eldridge *et al*., 2017</u>; <u>Fisch *et al*., 2019</u>; <u>Goddard *et al*., 2019</u>; <u>Mishra *et al*, 2023</u>; <u>Sanchez-Garrido *et al*, 2018</u>). Briefly, retroviral (with pMX backbones) and second-generation lentiviral packaging (with pTet backbones), pCMV-MMLV-Gag-Pol and pHIV-1266 were used, respectively, in combination with the pseudotyping plasmid pCMV-VSV-G-Env. HEK293E cells were seeded in 24-well format at a density of 1 x 10^5^ cells/well in 1 mL complete DMEM supplemented with 10 mM HEPES pH 7.5, and transfected with a total of 1 µg plasmid DNA per well using Lipofectamine 2000 in a 1:2.5 ratio (w:v) relative to DNA. For retroviral packaging, a 5:4:1 ratio of plasmid-of-interest:Gag-Pol:VSV-G was used. For Lentiviral packaging, a 3:2:1 ratio of plasmid-of-interest:HIV-1266:VSV-G was used. Virus-containing supernatants harvested 48 h post transfection, HEPES (pH 7.5) was added to 10 mM final concentration (to stabilise the pH of cell supernatants), and filter sterilised through 0.45 µm low-binding filters (Pall Life Sciences). For transductions, 250-400 µL of virus-containing supernatant was added to recipient cells in 24-well format. At 48 h post-transduction, the appropriate selection antibiotic was added and cells maintained in it until stable pools were obtained. If required, antibiotic-selected pools were sorted via FACS using an ARIA III (BD Bioscience) for uniform target gene expression.

### Immunofluorescence staining, image acquisition and processing

Cells were plated on glass cover slips in 24-well format and infected as described above. At specified timepoints post-infection, cover slips were washed twice with serum-free DMEM, and fixed using freshly prepared PBS containing 4 % PFA for 20 min at room temperature (RT, ∼20 °C). Cover slips were then washed twice with filter-sterilised PBS, and residual PFA was quenched using 50 mM NH4Cl prepared in PBS for 20 min at RT. Cover slips were washed twice with PBS, and, unless otherwise stated, permeabilised using Permeabilisation Buffer (PBS containing 0.3 % Triton X-100) for 3 min at RT, washed three times in PBS, and blocked in Blocking Buffer (PBS containing 5 % NDS) for 1 h before staining. Primary antibodies (***Reagents and Tools Table***) were diluted as per the manufacturer guidelines in Blocking Buffer + 0.1 % saponin and cover slips were inverted onto primary antibody solutions and incubated in a humidified chamber for 1 h at RT or 4 °C overnight. Cover slips were then washed three times in PBS in 24-well plates and inverted onto drops containing secondary antibodies (***Reagents and Tools Table***), phalloidin-Alexa Fluor-568 or phalloidin-Alexa Fluor-488 (Invitrogen), and Hoechst dye as appropriate in Blocking buffer for 1 h at RT in a humidified chamber. Cover slips were washed three times in filtered PBS, and mounted onto microscopy slides using ProLong Diamond Antifade Mountant (Fisher Scientific). Slides were allowed to cure for 24 h protected from light. In some experiments, imaging was performed in 96-well plates, and the same process was followed and samples imaged immediately following staining steps.

Images were acquired on a Zeiss CellDiscoverer 7 epifluorescence microscope fitted with an Axiocam 504 mono camera and a Colibri.2 light source. Samples were imaged at 100x magnification using a PApo 50x/1.2 water autoimmersion lens along with an afocal magnification lens with a range of 0.5-2x. The following LEDs were used: 385 / 470 / 520 / 567 / 590 / 625 nm along with Zeiss 90 HE, 91 HE and 92 HE filter sets. Images were processed using Zen Blue (Zeiss), and deconvolution and montages of pseudocoloured images were prepared using ImageJ (Fiji) software (<u>Schindelin *et al*, 2012</u>). Pearson’s Correlation Coefficient was calculated using the ImageJ plugin Coloc2 (Fiji) on regions of interest using original images (<u>Dunn *et al*, 2011</u>). Integrated pixel density was measured using Fiji. Deconvolution used theoretical point spread functions (PSFs) matching image acquisition conditions and were generated using the Richards-Wolf method using PSF Generator ImageJ plugin (<u>Kirshner *et al*, 2013</u>). DeconvolutionLab2 ImageJ plugin with the Richardson-Lucy method with 10-25 iterations and appropriate PSFs were used for deconvolution (<u>Sage *et al*, 2017</u>).

### Generation of human colonic epithelial organoid line

Three colonic mucosal biopsies from the sigmoid colon were obtained from a non-IBD control participant via lower gastrointestinal endoscopy, immediately placed in sterile PBS, and processed the same day for colonic epithelial organoid culture. Biopsies were washed three times in cold sterile PBS by gentle pipetting with a Pasteur pipette. Biopsies were then incubated in PBS containing 0.5 M EDTA and rotated at 4 °C for 40 min on a roller shaker. Following incubation, EDTA was removed by washing three times with cold PBS. Crypts were released by vigorous pipetting in 3 mL fresh PBS. The supernatant containing liberated crypts was collected into a fresh tube pre-coated with 0.1 % BSA in PBS. This step was repeated twice more to maximise yield. Collected fractions were pooled and centrifuged at 600 x*g* for 5 min at 4 °C. The pelleted crypts were resuspended in cold Matrigel ExtraCellular Matrix and 20–25 µL domes (two domes per well) were seeded into each well of a pre-warmed 24-well plate. Plates were incubated at 37 °C for 15-20 min to allow Matrigel to solidify before adding 500 µL of IntestiCult™ organoid growth media supplemented with 10 µM Rho Kinase inhibitor Y-27632 and 100 µg/mL Primocin per well. Organoids were grown at 37 °C and 5 % CO2 in a humidified incubator. Media was replaced the day after seeding and subsequently every 2-3 days.

### Passaging of colonic organoids

Organoids were passaged every 7–10 days. Culture medium was aspirated and ice-cold Gentle Cell Dissociation Reagent was added to each well to disrupt Matrigel domes. Gels were mechanically broken up and transferred into a pre-coated 15 mL tube on ice. Wells were rinsed with additional cold GCDR to recover residual Matrigel and organoids. Tubes were incubated on ice for 15-20 min to allow the Matrigel to dissolve, then centrifuged at 550 x*g* for 5 min at 4 °C. Supernatant was carefully removed. Pellets were resuspended in cold GCDR and dissociated by adding TrypLE Express. The suspension was pipetted up and down 2-3 times with a BSA-coated tip and incubated at 37 °C for 2-3 min. The reaction was stopped by diluting the cell suspension with cold PBS. Organoids were mechanically fragmented further by vigorous until the suspension appeared cloudy and fragments were mostly single cells or small clusters (checked microscopically). The suspension was centrifuged again, and the pellet was resuspended in fresh Matrigel. The cells were seeded and cultured as described above.

### Preparing organoid monolayers for infection and imaging

Human organoids were cultured as above in 40 μl drops of a 1:1 mixture of Matrigel and medium. Organoids from 12 drops were harvested into 1 mL of Gentle Cell Dissociation Reagent per drop and transferred to a tube precoated with 1% BSA, and kept on ice for 30 min. They were then pooled and 2 mL of Advanced DMEM:F12 medium was added to the pooled cells. The organoids were pipetted twice and centrifuged at 300 x*g* for 5 min. The medium was removed and 1 mL of TrypLE added to the pellet. Organoids were resuspended and incubated with frequent pipetting at 37 °C until single cells were visible under the microscope. DMEM:F12 (10 mL) was then added to the cells and the suspension centrifuged at 300 x*g* for 5 min. The pellet was resuspended in 1.2 mL of OGM and 100 μL added to one well of a 96-well plate precoated with Matrigel. For precoating, 96w plates were treated with 1:50 dilution of Matrigel prepared in DPBS for 1 h, the solution removed and cells were plated directly on coated wells. Cells were allowed to attach and grow until cells became confluent. The medium was then changed to Organoid Differentiation Medium and culturing continued for a further 4 days to allow differentiation prior to experiments.

### Triggering FcγR-Tir signalling with hIgG-coated beads

HeLa cells encoding FcγR-Tir fused to a C-terminal mCherry2 or MYC tag were seeded on glass coverslips in 24-well plates at a density of 8 x 10^4^ cells/well for 48 h. Cells were treated with 10 ng.mL-^1^ IFNγ for 16 h before treatment with beads. Polystyrene beads (3.54 µm diameter, Spherotech) were pelleted at 5,000 x*g* for 1 min, washed in 70 % ethanol and thrice in sterile PBS. For coating, beads were suspended in PBS containing either 50 µg.mL^-1^ of human IgG or 50 µg.mL^-1^ BSA and incubated for 18 h on a rotating wheel (10 rpm) at 4 °C. The beads were then washed three times in sterile PBS. Beads were counted and added to cells (at a density of 5 beads/cell) followed by centrifugation of the culture plate at 300 x*g* for 5 minutes. Incubation was carried out at 37°C with 5 % CO_2_ for 15 mins, followed by washing with serum-free DMEM, and replacing media with OptiMEM (supplemented with 1 mM sodium pyruvate). After 3 h, cells were washed twice with ice-cold PBS and fixed with 4 % PFA and samples prepared for immunofluorescence as described above. The integrated density of caspase-4 was calculated using ImageJ (Fiji) software by thresholding against actin-rich structures stained with phalloidin (MaxEntropy, Fiji), expanding a region of interest and quantifying total intensity for caspase-4 staining within the region.

### Data handling and statistics

For all quantitative experiments, a single mean was calculated from two or three technical replicates (i.e. replicate wells in a 96-well plate for PI-uptake assays). Data were excluded if an entire experiment failed technically (e.g., low infection level or <10 % cell death). Experiments were repeated using distinct passages of cells on different days and independent cultures of bacteria to generate biologically and statistically independent data (represented by n in figure legends). Data were analysed using R (v.4.0 or higher, grafify package (<u>Shenoy, 2021</u>)) or GraphPad Prism (v.8.0 or higher) (***Reagents and Tools Table***). For bar charts, mean and standard deviation (SD) are displayed. For box plots, median and interquartile range (IQR) are indicated as defined in the figure legend, with boxes indicating IQR, whiskers indicating 1.5x IQR and the line indicating the median. Data plotting using grafify and ggplot2 (<u>Wickham, 2016</u>) packages in R. Blinding was not used, samples sizes were not pre-determined and treatments were randomly assigned mice.

Statistical comparisons used mixed effects linear models fitted without or with random intercepts (experimental blocks) and analysed using the grafify R package. Residual distributions were analysed to ensure normal distribution when using parametric analyses, and log or logit transformations were used where necessary. Post-hoc comparisons used FDR adjustment (α = 0.05) used to correct *P* values for multiple comparisons.

### Ethics declarations

All animal experiments complied with the Animals Scientific Procedures Act 1986 and U.K. Home Office guidelines and were approved by the local ethical review committee. Experiments were designed in agreement with the ARRIVE guidelines (<u>Kilkenny *et al*, 2010</u>) for the reporting and execution of animal experiments, including sample randomization and blinding. This study complies with all relevant ethical regulations for research involving human participants. The use of human colonic samples to generate organoids was conducted under the THAMES-IBD study protocol sponsored by Imperial College London, which was reviewed and approved by the Health Research Authority Sheffield Research Ethics Committee (IRAS ID: 290708). Informed consent was obtained from all participants in accordance with the approved protocol. All samples were pseudonymised prior to analysis.

## Data availability

Primary data in figures are uploaded at Bioimage Archive and accessible here: https://www.ebi.ac.uk/biostudies/bioimages/studies/S-BIAD3354. The Expanded view data are available here:

## Acknowledgements

Wouter W Kallemeijn is sadly deceased; this manuscript is dedicated to his memory. Credit statement: Investigation – DB, IC, DC, JCNW, AP, PB, MS, MG, QZ, ARS; Formal analysis – DB, IC, DC, JCNW, AP, PB, MS, QZ, ARS; Resources –MG, WWK, SK, DP, JPT, DD, TK, GF; Methodology – DB, WWK, EWT, SK, DP, JPT, DD, TK, SSV, ARS; Software – DB, IC, ARS; Writing – original draft – DB, ARS; Writing – review & editing – DB, IC, DC, PB, AMT, EMF, EWT, SSV, GF, ARS; Conceptualisation – AMT, EMF, SSV, EWT, GF, ARS; Funding acquisition – DC, SSV, EWT, EMF, ARS. Funding sources – UK Medical Research Council (MR/V030930/1 to ARS, EMF, EWT), UK Commonwealth Scholarship Commission (INCN-2024-75 to DC, SSV and ARS), Council of Scientific & Industrial Research, India (to DC), National Institutes of Health, USA (1R01AI169618-01A1 to SSV and ARS), Cancer Research UK, with support from the Engineering & Physical Sciences Research Council, UK (DRCNPG-Nov21\100001 to EWT). The Francis Crick Institute receives its core funding from Cancer Research UK (FC001057, 361 FC001002), the UK Medical Research Council (MRC; FC001057, FC001002), and the Wellcome Trust (FC001057, FC001002). SK and DP were supported by the National Institute for Health Research (NIHR) Imperial Biomedical Research Centre (BRC). JPT is supported by the Chain-Florey Clinical PhD Fellowship jointly funded by the NIHR Imperial Biomedical Research Centre (BRC) and the UKRI Medical Research Council (MRC) Laboratory of Medical Sciences (LMS). We thank the Facility for Light Microscopy (FILM) and Flow Cytometry Facility at Imperial College London. We thank Xinru Li, Abigail Iles, Anne Schumacher and Sarah Hassan for helpful discussions, Philippa Goddard for help with initial FcγR-Tir experiments, and Julia Sanchez-Garrido and Abigail Clements for comments on the manuscript draft. We thank Dr P Susmitha Karuna Sree and Prof V Balaji of the Dept of Microbiology, Christian Medical College, Vellore, Tamil Nadu, India for sharing the EPEC O119:H6 isolate. The authors would like to express their acknowledgements to Nick Powell and Aamir Saifuddin for establishing the THAMES-IBD cohort that provided samples for the organoid generation. The views expressed are those of the authors and not necessarily those of the NIHR or the UK Department of Health and Social Care.

## Reagents and tools table

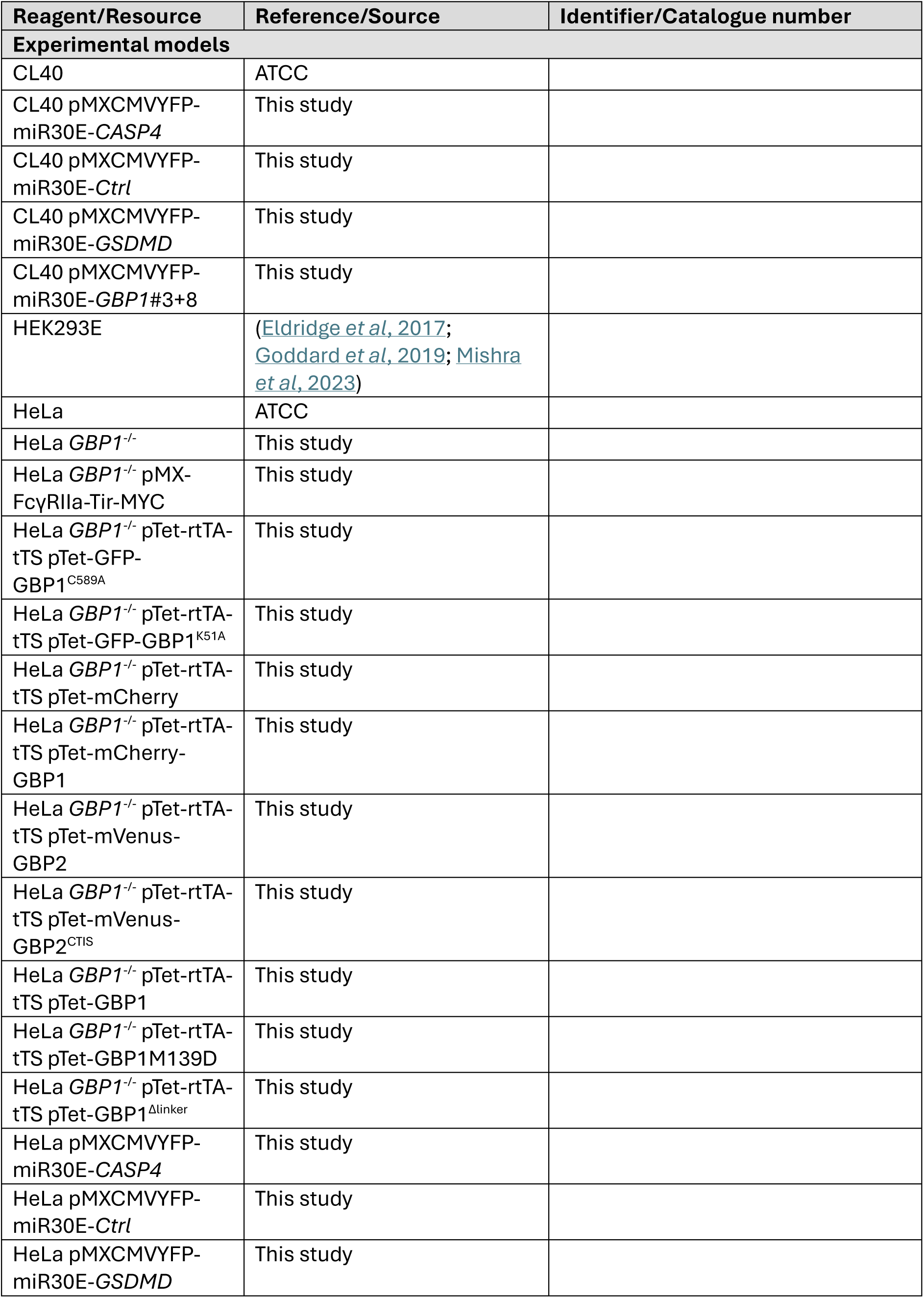
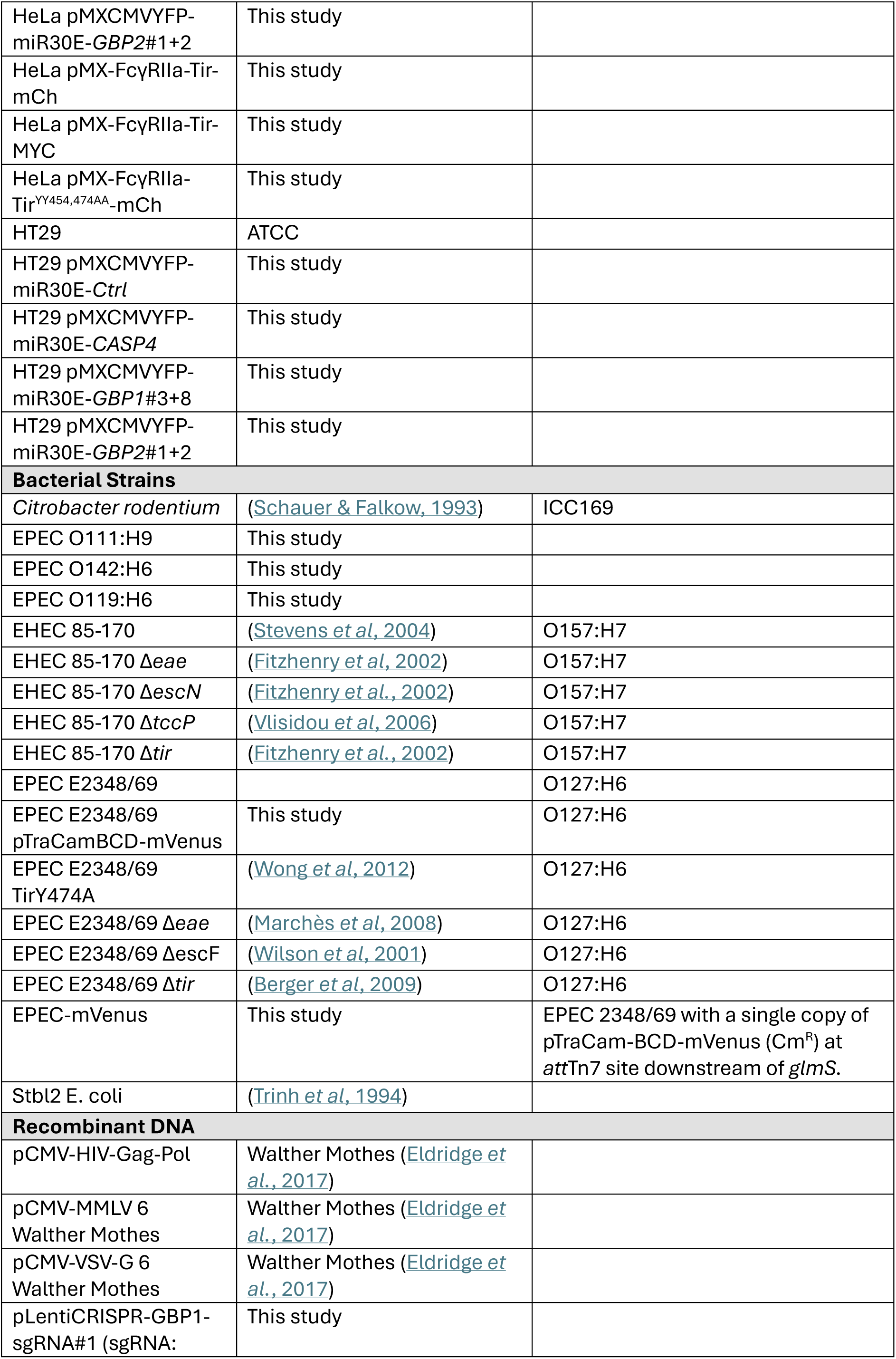
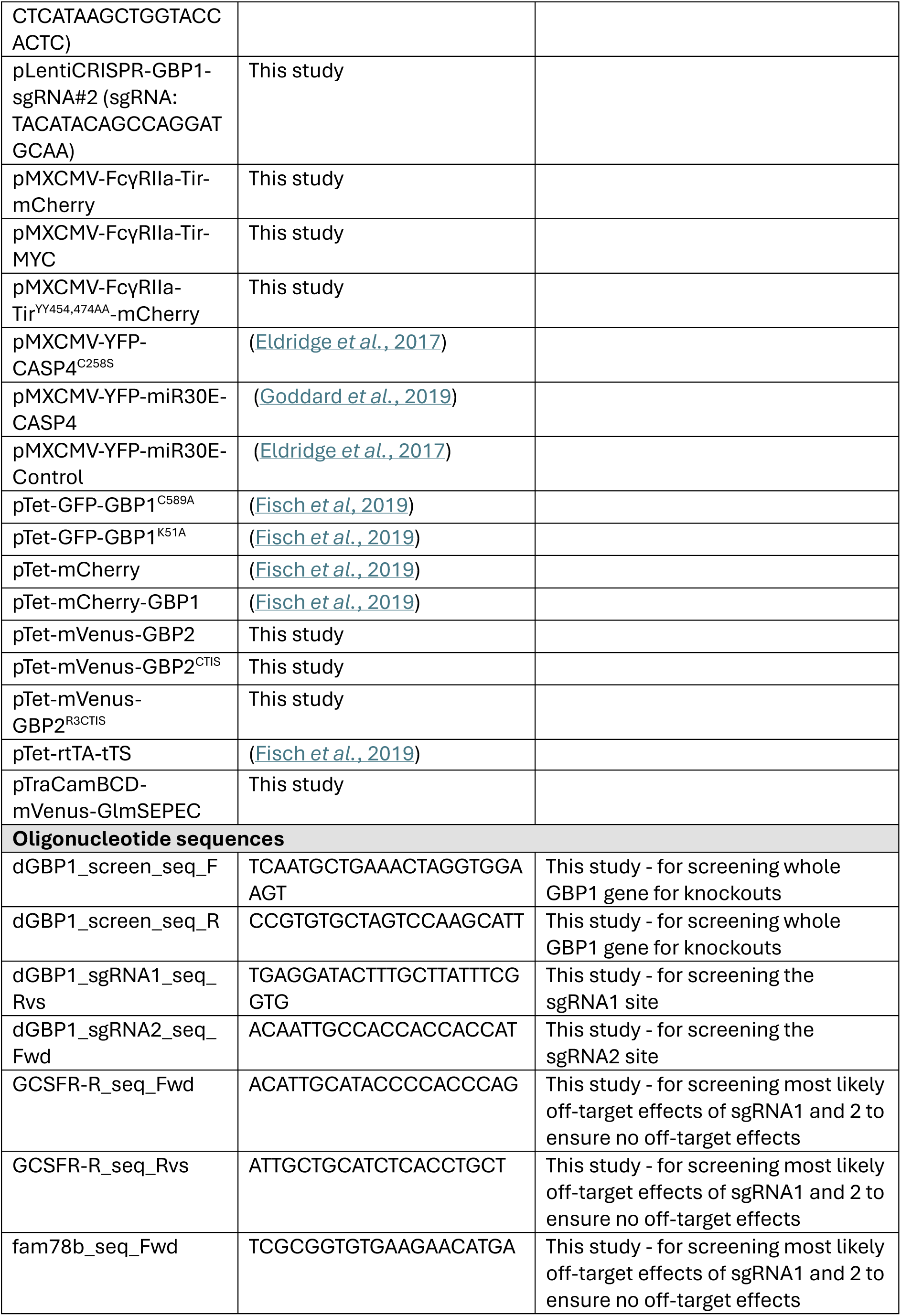
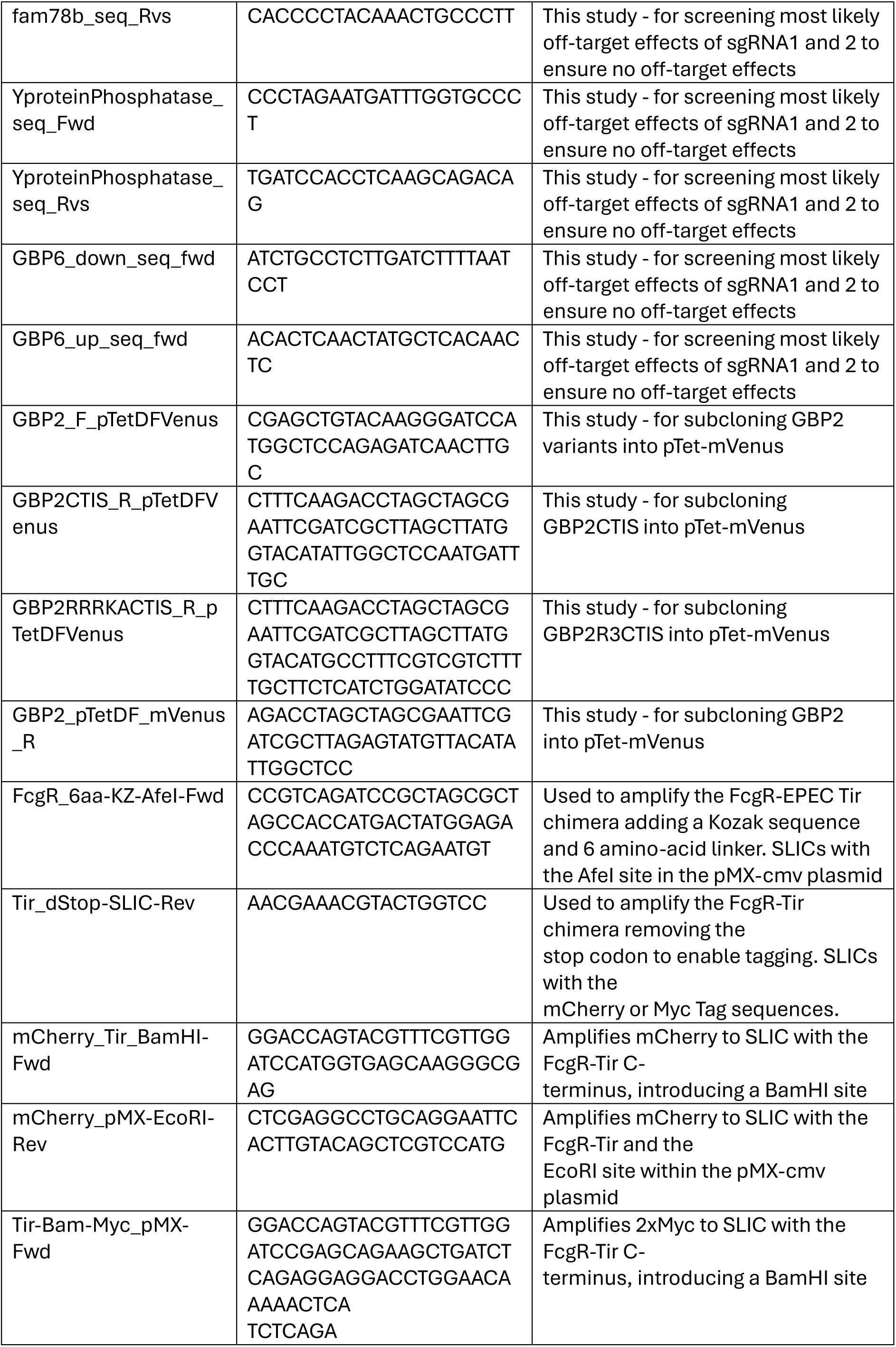
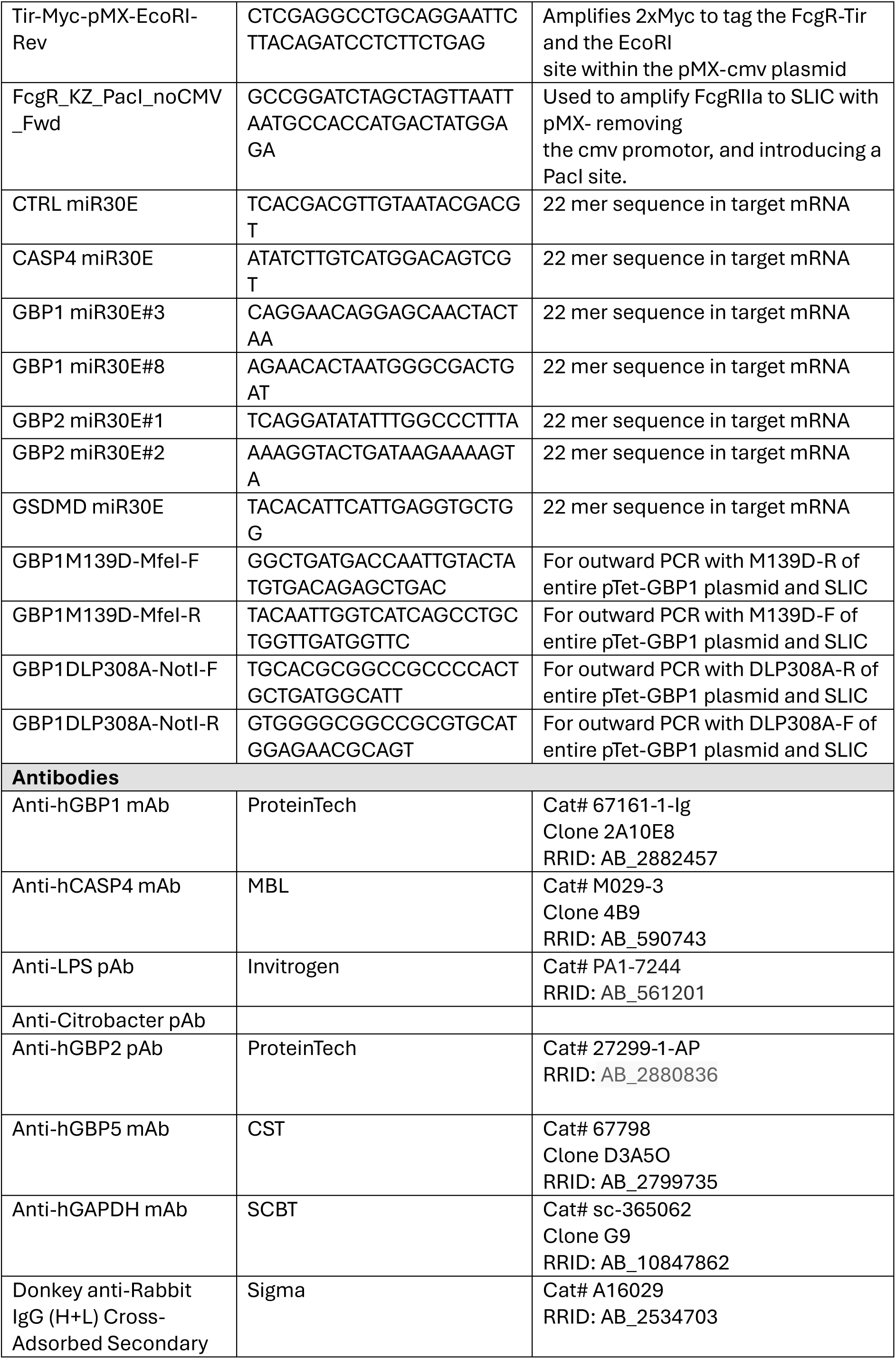
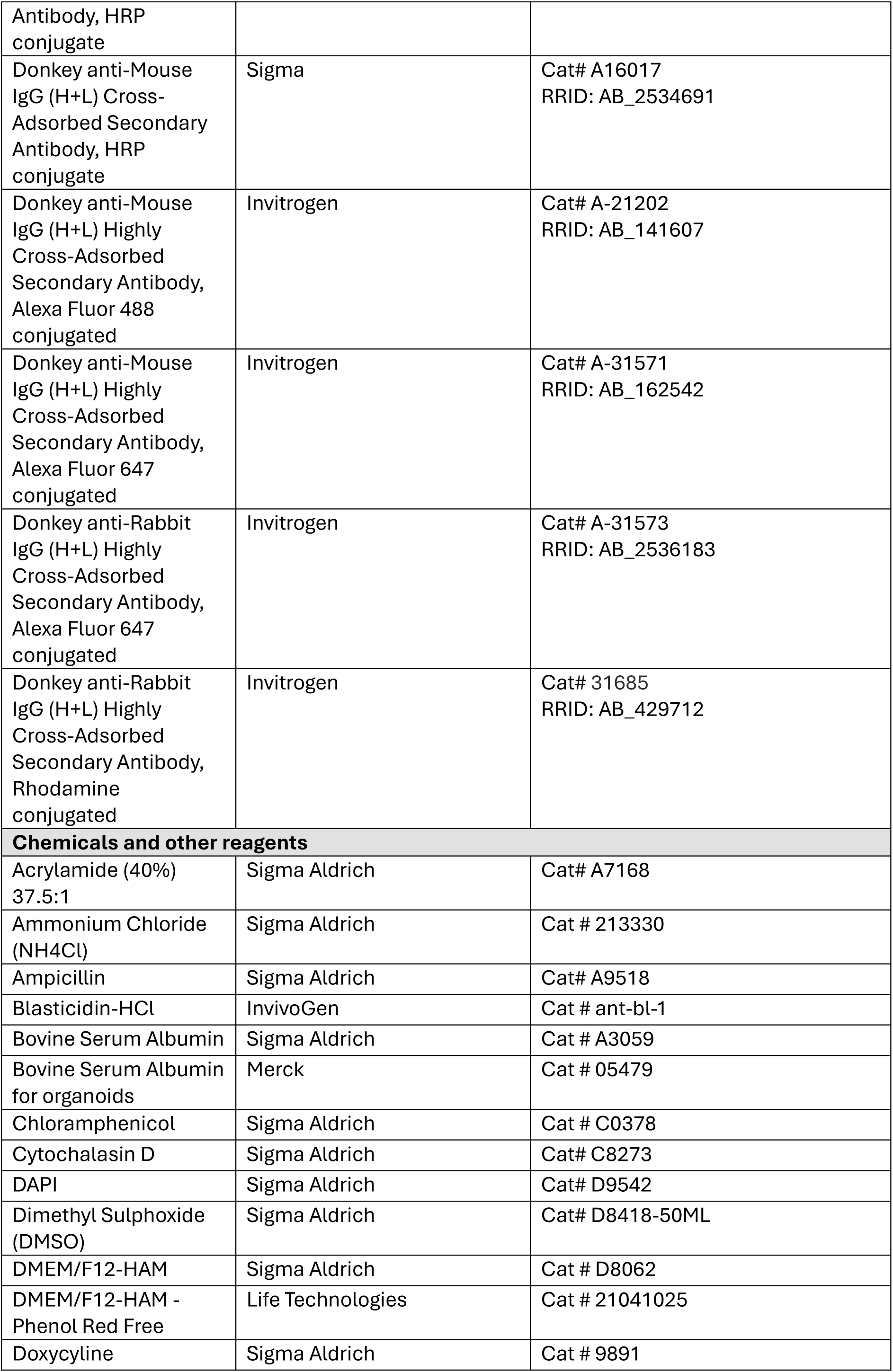
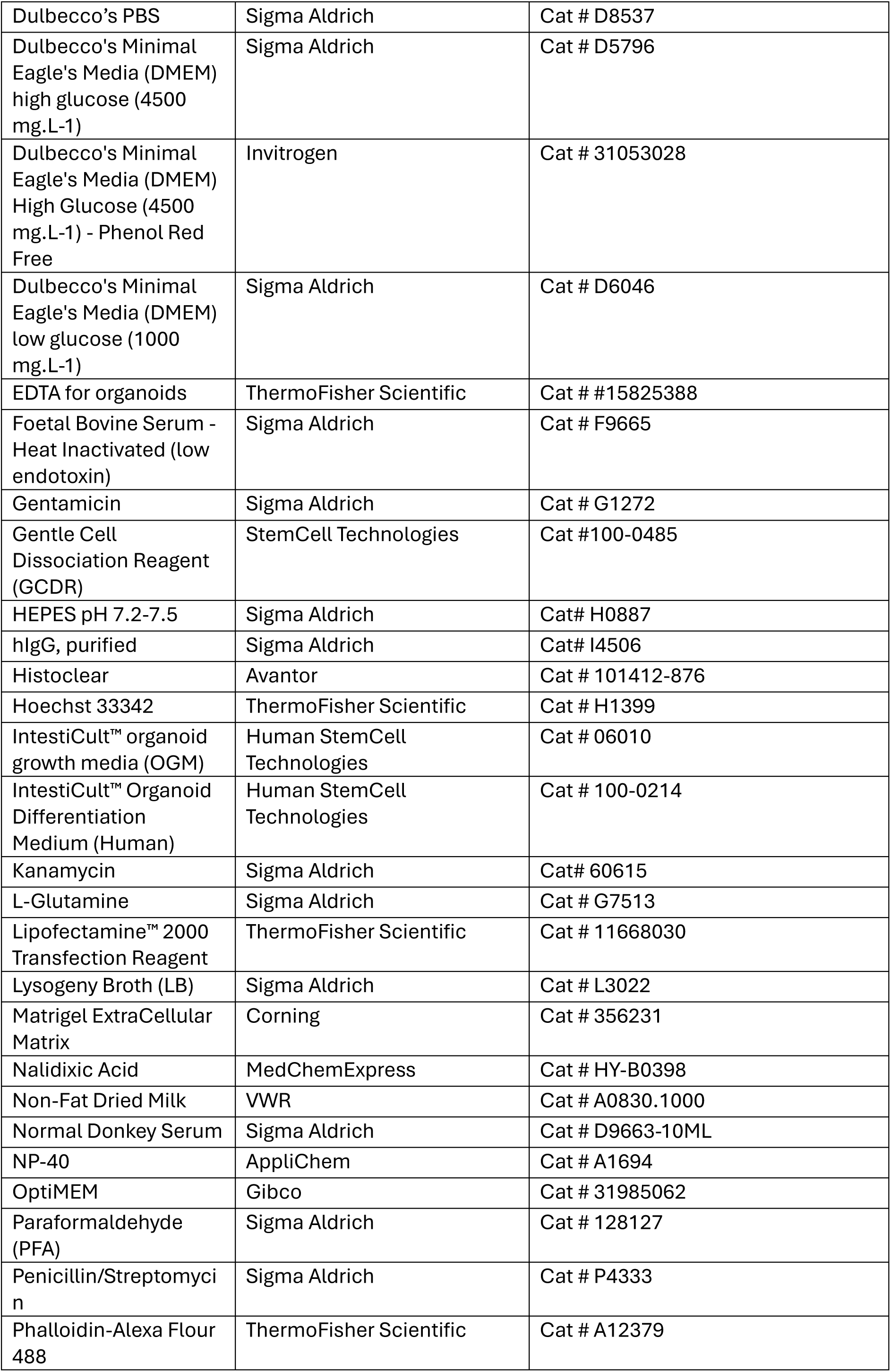
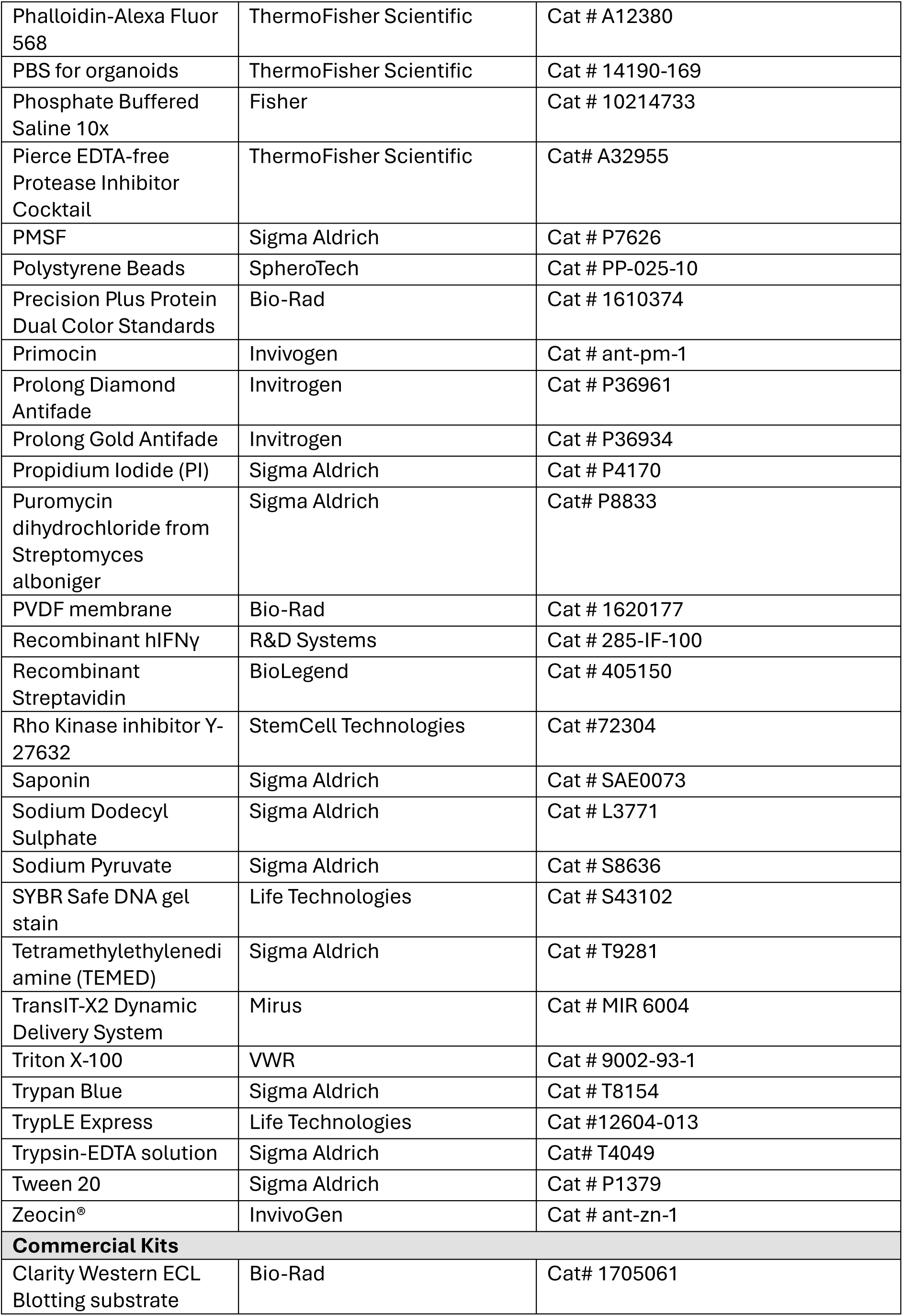
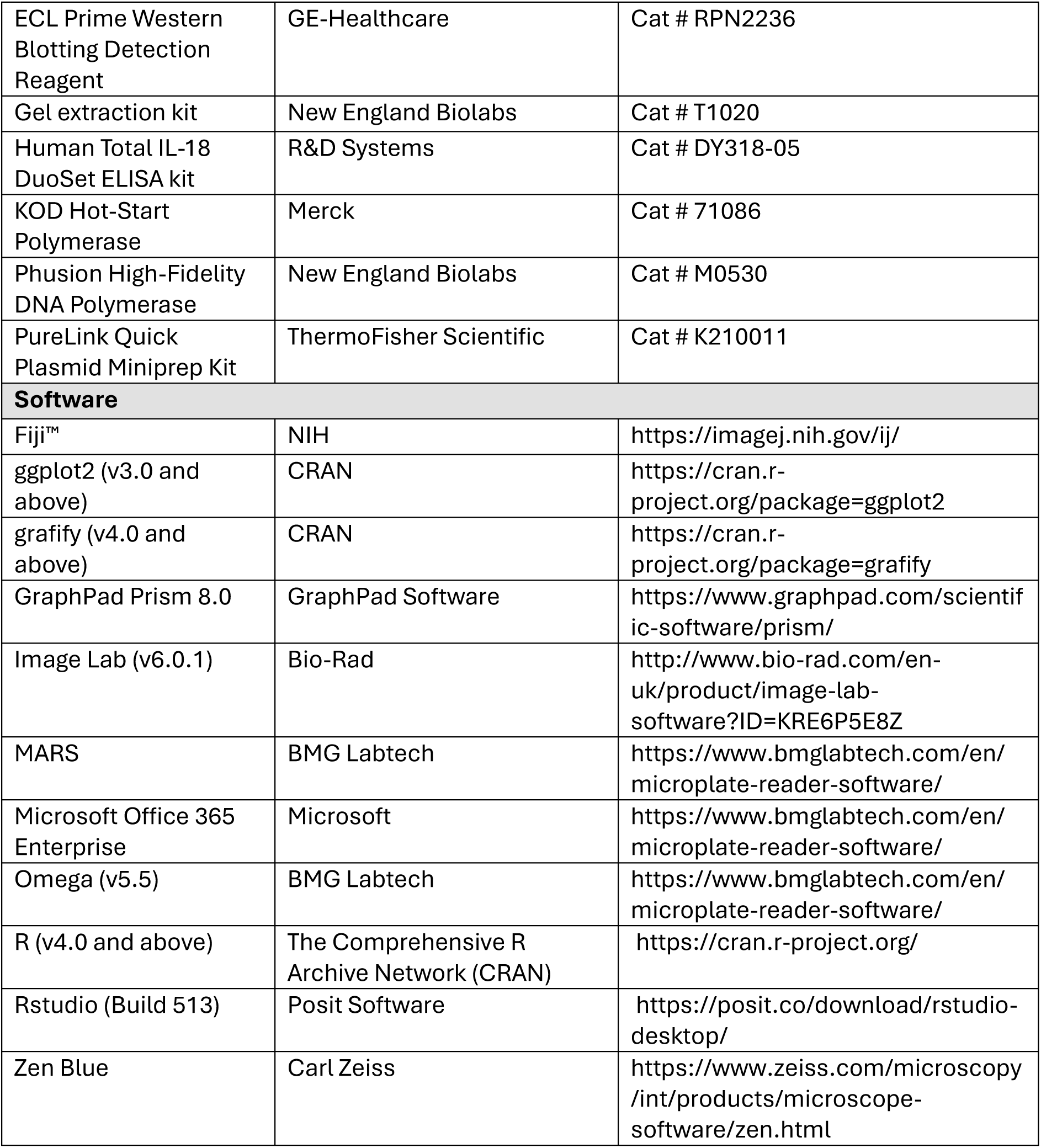

