## Supplementary figures for "GBP1 recruitment to actin-rich pedestals induced by extracellular Gram-negative bacteria promotes pyroptosis"

### Supplementary Figures EV1-EV5

**Bennison et al**
**Figure EV1**

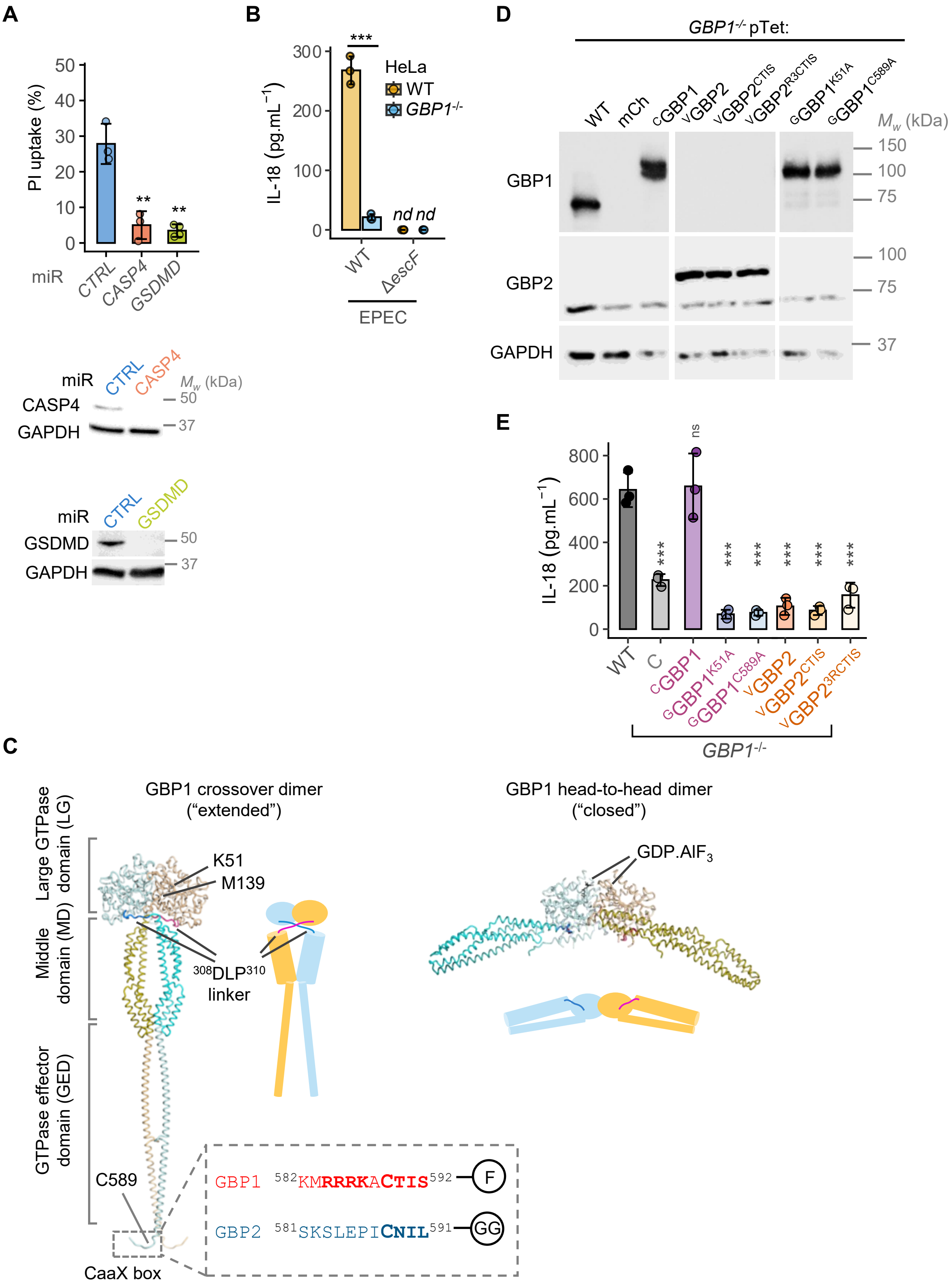

**Figure EV1. GBP1 is recruited to actin-rich pedestals of attaching and effacing (A/E) pathogens.**

(A) Graph showing percentage pyroptotic cell death (top) and representative immunoblots (below) of HeLa cells stably expressing non-targeting (negative control; CTRL) or miR30E targeting caspase-4 (*CASP4*) or gasdermin D (*GSDMD*). Graph (top) shows percentage propidium iodide dye uptake assays in the indicated cells infected with EPEC for 6 h. Cells were primed with IFNγ (10 $ng.mL^{-1}$) 16 h before infection or preparation of cell lysates for western blots. Mean ± SD error bars with symbols representing data from n = 3 independent experiments are shown. ** $P < 0.01$, two-tailed $P$ values for comparisons of CTRL and the indicated miRNA-expressing cells from mixed effects ANOVAs ($P = 0.0037$ for the two indicated comparisons). Representative immunoblots (below) of caspase-4, GSDMD and GAPDH (loading control) from n = 3 independent experiments.

(B) ELISA quantification of IL-18 from supernatants of IFNγ-primed HeLa cells of the indicated genotypes infected with wild-type (WT) or Δ*escF* mutant of EPEC for 6 h. Mean ± SD error bars with symbols representing data from n = 3 independent experiments are shown. *** $P < 0.001$ ($P = 2.6e-07$) is a two-tailed $P$ value for the indicated comparisons from mixed effects ANOVAs. *nd*, not detected.

(C) Ribbon structures of "extended" or "open" crossover dimer of GBP1 on left (adapted from PDB: 8R1A), and "closed" or "safety pin" dimer (modelled on PDB: 2B92 LG dimer bound to GDP.$AlF_3$) shown on right. GBP1 monomers coloured pastel shades of cyan and wheat, respectively. Residues K51, C589, M139, $^{308}$DLP$^{310}$ linker, GDP.$AlF_3$ and the farnesyl group are highlighted. C-terminal sequences for GBP1 and GBP2 are described with CaaX-box motifs and the GBP1 polybasic RRR motif shown in bold. The prenylation state of the protein is indicated with F or GG (farnesyl and geranylgeranyl, respectively).

(D) Representative immunoblots showing the expression of the indicated tetracycline (Tet)-controlled WT GBP1 or the indicated variants in *GBP1$^{-/-}$* cells. Cells were stably transduced with expression plasmids for the indicated proteins, treated with IFNγ (10 $ng.mL^{-1}$) and doxycycline (200 $ng.mL^{-1}$) for 16 h and cell lysates prepared for western blots. Data are representative of n = 2 independent experiments. Images shown are cropped from the same membrane at the same exposure to remove unnecessary lanes.

(E) ELISA quantification of IL-18 from supernatants of IFNγ- and doxycycline-treated wild-type (WT) or *GBP1$^{-/-}$* HeLa cells stably expressing mCherry2 (C), mCherry2-tagged WT GBP1 or K51A or C589A mutants, or mVenus-GBP2 or variants as indicated. Mean ± SD error bars with symbols representing data from n = 3 independent experiments are shown. *ns*, not significant; *** $P < 0.001$, two-tailed $P$ values for comparisons between WT cells and cells expressing the indicated GBP1 or GBP2 variants from mixed effects ANOVAs. Exact $P$ values as follows – C: 1.7e-06; cGBP1: 0.9282; $^{G}$GBP1$^{K51A}$: 5.5e-10; $^{G}$GBP1$^{C589A}$: 5.5e-10; $^{V}$GBP2: 1.8e-09; $^{V}$GBP2$^{CTIS}$: 7.8e-10; $^{V}$GBP2$^{3RCTIS}$: 3.2e-08.

**Bennison et al**
**Figure EV2**

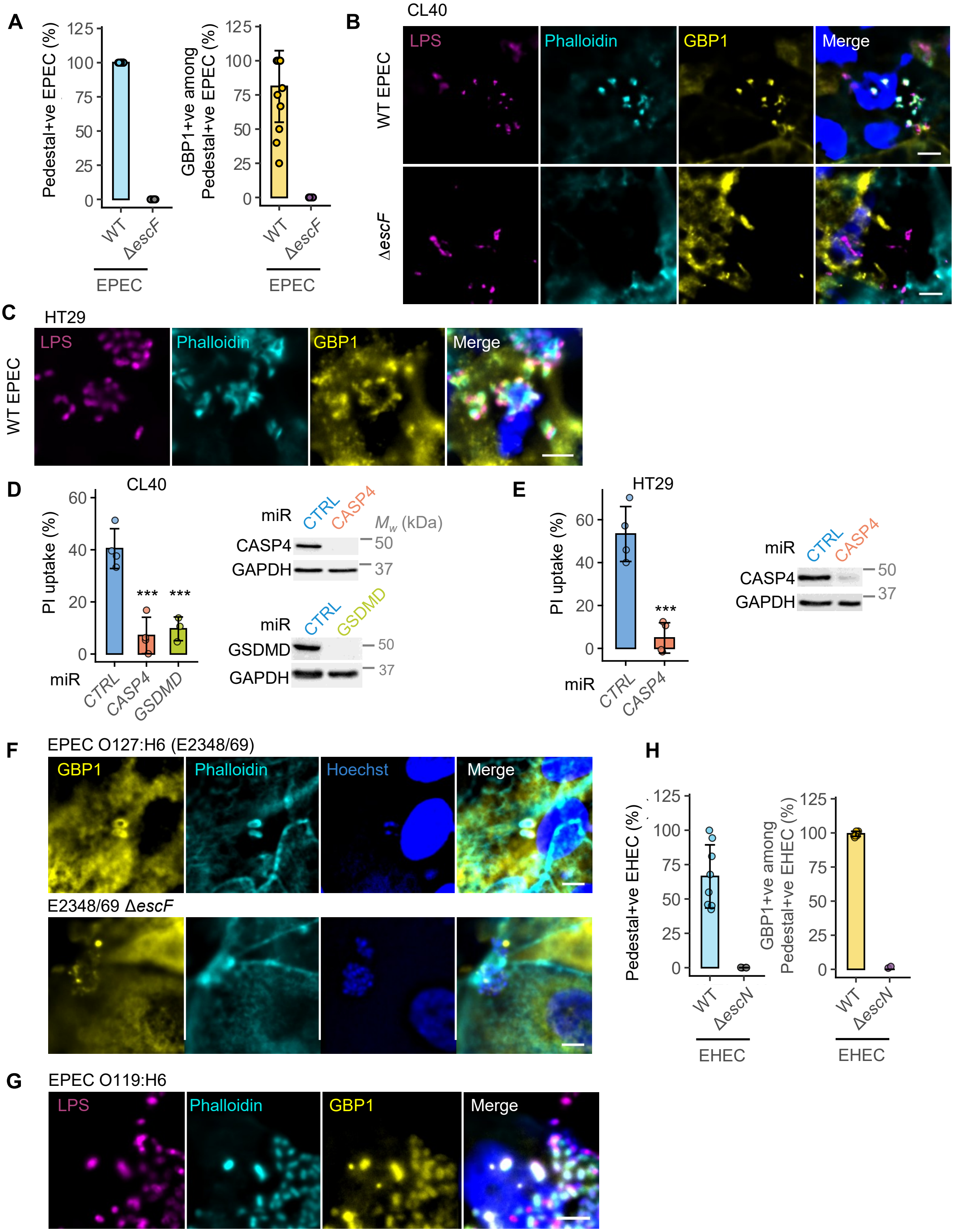

**Figure EV2. GBP1 is recruited to actin-rich attachment sites of extracellular EPEC.**

(A) Quantification of microcolonies of wild-type (WT) or Δ*escF* mutant of EPEC infected in IFNγ-primed HeLa cells staining positive for actin-rich pedestals observed with phalloidin staining (Pedestal+ve) or positive for both phalloidin and GBP1 (GBP1+ve among Pedestal+ve) at 2 h post-infection (also see images in Figure 1). Mean ± SD error bars with symbols representing individual fields of view from n = 3 independent experiments are shown.

(B) Representative immunofluorescence images of IFNγ-primed human CL40 colonic epithelial cells infected with WT EPEC or Δ*escF* mutant for 1 h. Cells were stained with anti-LPS and anti-GBP1 antibodies, phalloidin-Alex568 (to stain actin) and Hoechst (DNA dye; blue). Data are representative of n = 3 independent experiments. Scale bar, 5 μm.

(C) Representative immunofluorescence images of IFNγ-primed HT29 cells infected with EPEC E2348/69 for 3 h. Cells were stained with anti-GBP1 antibodies, phalloidin-Alex568 (to stain actin) and Hoechst (DNA dye; blue). Data from n = 2 independent experiments. Scale bar, 5 μm.

(D) Graph showing percentage pyroptotic cell death (left) at 6 h post-infection with EPEC and representative immunoblots (right) of CL40 cells stably expressing negative control (CTRL) or miR30E targeting caspase-4 (*CASP4*) or gasdermin D (*GSDMD*). Cells were primed with IFNγ (10 $ng.mL^{-1}$) 16 h before infection. Mean ± SD error bars with symbols representing data from n = 3 independent experiments are shown. *** $P < 0.001$, two-tailed *P* values for comparisons between CTRL and indicated miRNA-expressing cells from mixed effects ANOVAs ($P = 0.0008$ for the two indicated comparisons). Representative immunoblots (right) of caspase-4, GSDMD and GAPDH (loading control) from n = 3 independent experiments.

(E) Graph showing percentage pyroptotic cell death (left) at 12 h post-infection with EPEC and representative immunoblots (right) of HT29 cells stably expressing negative control (CTRL) or miR30E targeting caspase-4 (*CASP4*). Cells were primed with IFNγ (10 $ng.mL^{-1}$) 16 h before infection. Mean ± SD error bars with symbols representing data from n = 4 independent experiments are shown. *** $P < 0.001$, two-tailed *P* values for comparisons between CTRL and *CASP4*-miRNA-expressing cells from mixed effects ANOVAs are indicated ($P = 9.1e\text{-}07$). Representative immunoblots (right) of caspase-4 and GAPDH (loading control) from n = 3 independent experiments.

(F) Representative immunofluorescence images of IFNγ-primed 2D monolayers prepared from human colonic organoids infected with wild-type EPEC or Δ*escF* mutant for 2 h as indicated. Cells were stained with anti-GBP1 antibodies, phalloidin (to stain actin) and Hoechst (DNA dye; blue) as labelled. Scale bar, 5 μm.

(G) Representative immunofluorescence images of IFNγ-primed HeLa infected with the indicated strain of EPEC for 3 h. Cells were stained with anti-LPS and anti-GBP1 antibodies, phalloidin (to stain actin) and Hoechst (DNA dye; blue). Data are representative of n = 3 independent experiments. Scale bar, 5 μm.

(H) Quantification of wild-type (WT) or Δ*escN* mutant of EHEC infected in IFNγ-primed HeLa cells staining positive for actin-rich pedestals observed with phalloidin staining (Pedestal+ve) or positive for both phalloidin and GBP1 (GBP1+ve among Pedestal+ve) at 5 h post-infection (also see images in Figure 1). Mean ± SD error bars with symbols representing individual fields of view from n = 3 independent experiments are shown.

**Bennison et al**
**Figure EV3**

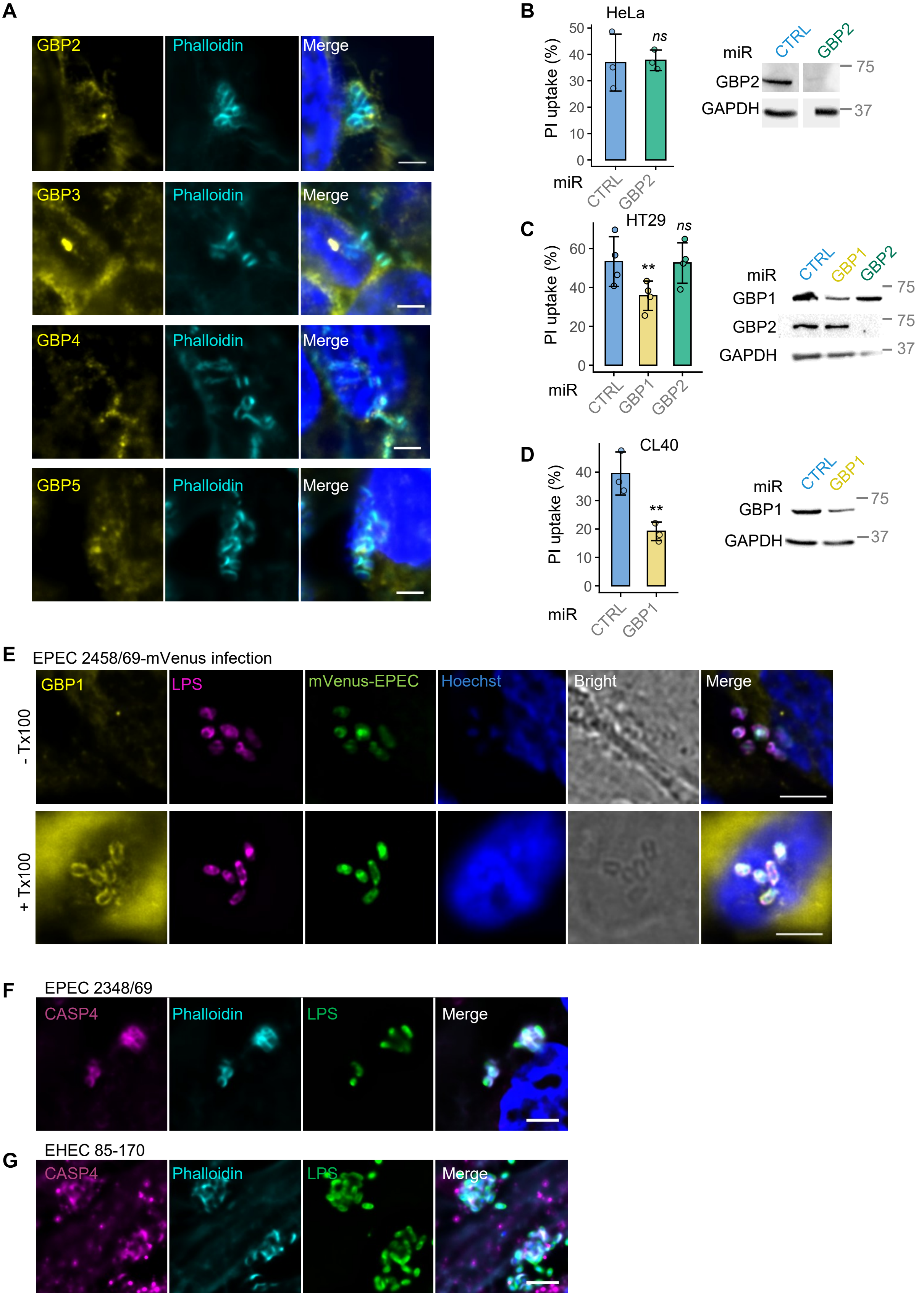

**Figure EV3. GBP1 is recruited to actin-rich attachment sites of extracellular EPEC.**

(A) Representative immunofluorescence images of IFNγ-primed and doxycyline-treated HeLa cells showing the expression of the indicated GBPs and infected with EPEC for 3 h. Cells were fixed and $^{FLAG}$GBP2, $^{FLAG}$GBP3, $^{MYC}$GBP4 and $^{FLAG}$GBP5 were strained with anti-FLAG or anti-MYC antibodies, phalloidin (to stain actin) and Hoechst (DNA dye; blue). Data are representative of n = 3 independent experiments. Scale bar 4 μm.

(B) Graph showing percentage pyroptotic cell death (left) at 6 h post-infection with EPEC and representative immunoblots (right) of HeLa cells stably expressing negative control (CTRL) or miR30E targeting *GBP2*. Cells were primed with IFNγ (10 ng.mL$^{-1}$) 16 h before infection. Mean ± SD error bars with symbols representing data from n = 3 independent experiments are shown. *ns*, not significant (*P = 0.8541*) for comparison between CTRL and GBP2 miRNA-expressing cells from mixed effects ANOVAs. Representative immunoblots (right) of GBP2 and GAPDH (loading control) from n = 3 independent experiments. Images show the same exposures of the same blots for GBP2 and GAPDH with irrelevant lanes removed.

(C) Graph showing percentage pyroptotic cell death (left) at 12 h post-infection with EPEC and representative immunoblots (right) of HT29 cells stably expressing negative control (CTRL) or miR30E targeting *GBP1* or *GBP2*. Cells were primed with IFNγ (10 ng.mL$^{-1}$) 16 h before infection. Mean ± SD error bars with symbols representing data from n = 4 independent experiments are shown. ** *P = 0.0014; ns*, not significant (*P = 0.8417*) are two-tailed *P* values for comparison of CTRL with the indicated miRNA-expressing cells from mixed effects ANOVAs. Representative immunoblots (right) of GBP1, GBP2 and GAPDH (loading control) from n = 4 independent experiments.

(D) Graph showing percentage pyroptotic cell death (left) at 6 h post-infection with EPEC and representative immunoblots (right) of CL40 cells stably expressing negative control (CTRL) or miR30E targeting *GBP1*. Cells were primed with IFNγ (10 ng.mL$^{-1}$) 16 h before infection. Mean ± SD error bars with symbols representing data from n = 3 independent experiments are shown. ** *P = 0.0079* is a two-tailed *P* values for comparisons between CTRL and GBP1 miRNA-expressing cells from mixed effects ANOVAs. Representative immunoblots (right) of GBP1 and GAPDH (loading control) from n = 3 independent experiments.

(E) Representative immunofluorescence images of IFNγ-primed HeLa cells infected with EPEC-mVenus for 1 h and then fixed and stained without and with permeabilization as labelled. Cells were fixed and either permeabilised for 3 min with 0.3 % Triton X100 or not prior to staining with anti-LPS and anti-GBP1 antibodies, and Hoechst (DNA dye; blue). Data are representative of n = 3 independent experiments. Scale bar, 5 μm.

(F) – (G) Representative immunofluorescence images of IFNγ-primed HeLa cells infected with EPEC 2348/69 for 2 h (A) or EHEC 85-170 for 5 h (B) showing endogenous caspase-4 trafficking to actin-rich pedestals induced by bacteria. Cells were stained with anti-LPS (to stain bacteria) and anti-caspase-4 antibodies, phalloidin (to stain actin), and Hoechst (DNA dye; blue). Data are representative of n = 2 independent experiments. Scale bar, 5 μm.

**Bennison et al**
**Figure EV4**

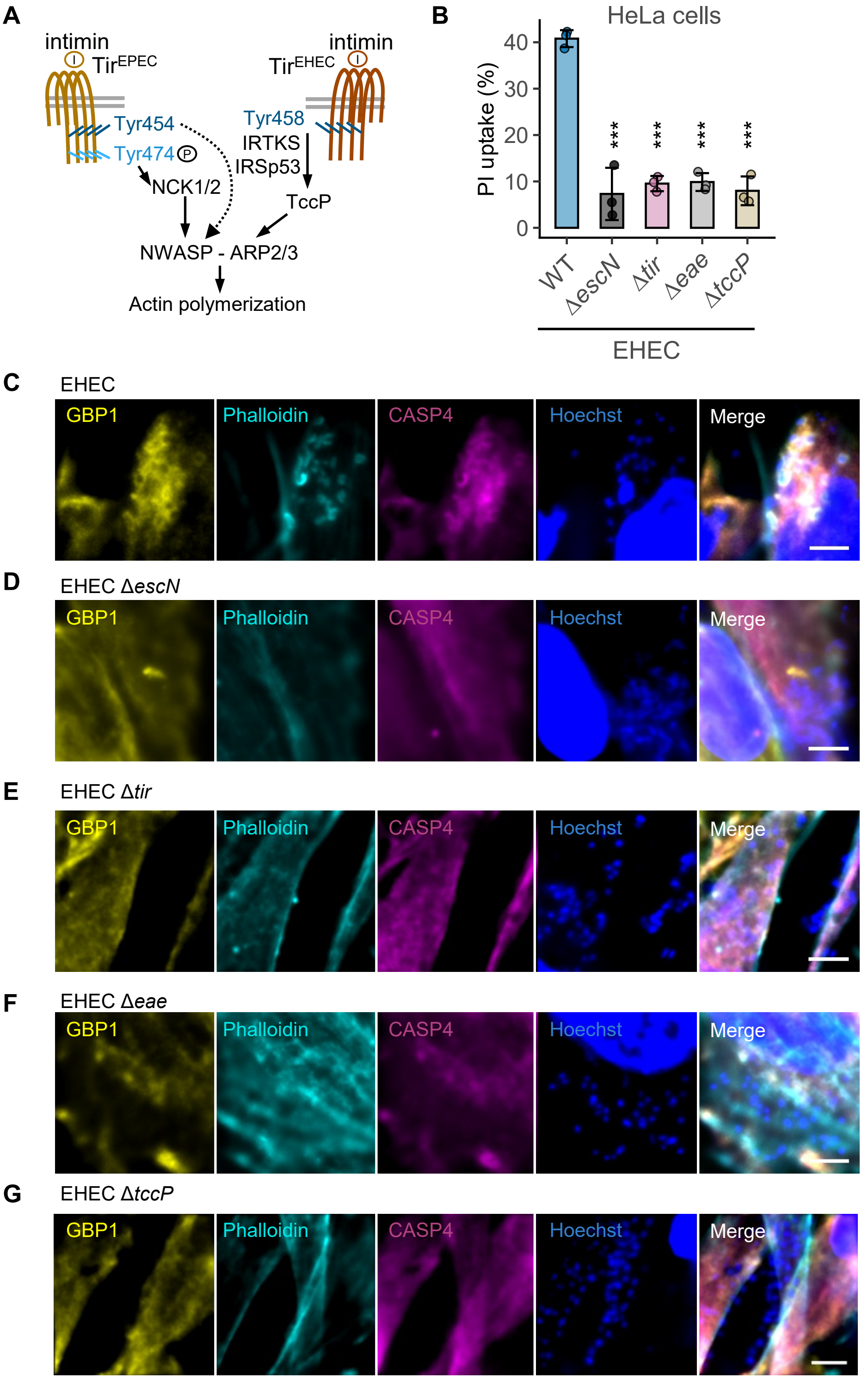

**Figure EV4. Tir-intimin signalling is essential for GBP1-caspase-4 recruitment to actin-rich pedestals of EHEC.**

(A) A schematic showing the distinct pathways of actin polymerisation driven by $Tir^{EPEC}$ and $Tir^{EHEC}$. $Tir^{EPEC}$ mainly relies on Y474 tyrosine phosphorylation, NCK recruitment and N-WASP-ARP2/3—dependent actin polymerisation. $Tir^{EHEC}$ uses Y458 to recruit IRTKS/IRSp53 which bridge the interaction between Tir, TccP and N-WASP-ARP2/3. The Y454 residue in $Tir^{EPEC}$ can also recruit IRTKS/IRSp53, but this does not lead to actin-rich structures due to the lack of TccP in EPEC 2348/69 (shown by dotted arrow)

(B) Percentage pyroptotic cell death as measured by propidium iodide dye uptake assays of IFNγ-primed HeLa cells infected with wild-type EHEC (WT) or the indicated mutants for 8 h. Mean ± SD error bars with symbols representing data from n = 3 independent experiments are shown. *** $P < 0.001$, two-tailed $P$ values for comparisons of the indicated mutant strains with the WT from mixed effects ANOVAs. $P$ values as follows – Δ*escN*: 1.6e-08; Δ*tir*: 1.9e-08; Δ*eae*: 1.9e-08; Δ*tccP*: 1.6e-08.

(C) – (G) Representative immunofluorescence images of IFNγ-primed HeLa cells expressing YFP-$Caspase4^{C285S}$ infected with wild-type EHEC or the mutants for 5 h. Cells were stained with anti-GBP1 antibodies, phalloidin-Alex568 (to stain actin) and Hoechst (DNA dye; blue). Data from n = 3 independent experiments. Scale bar, 5 μm.

**Bennison et al**
**Figure EV5**

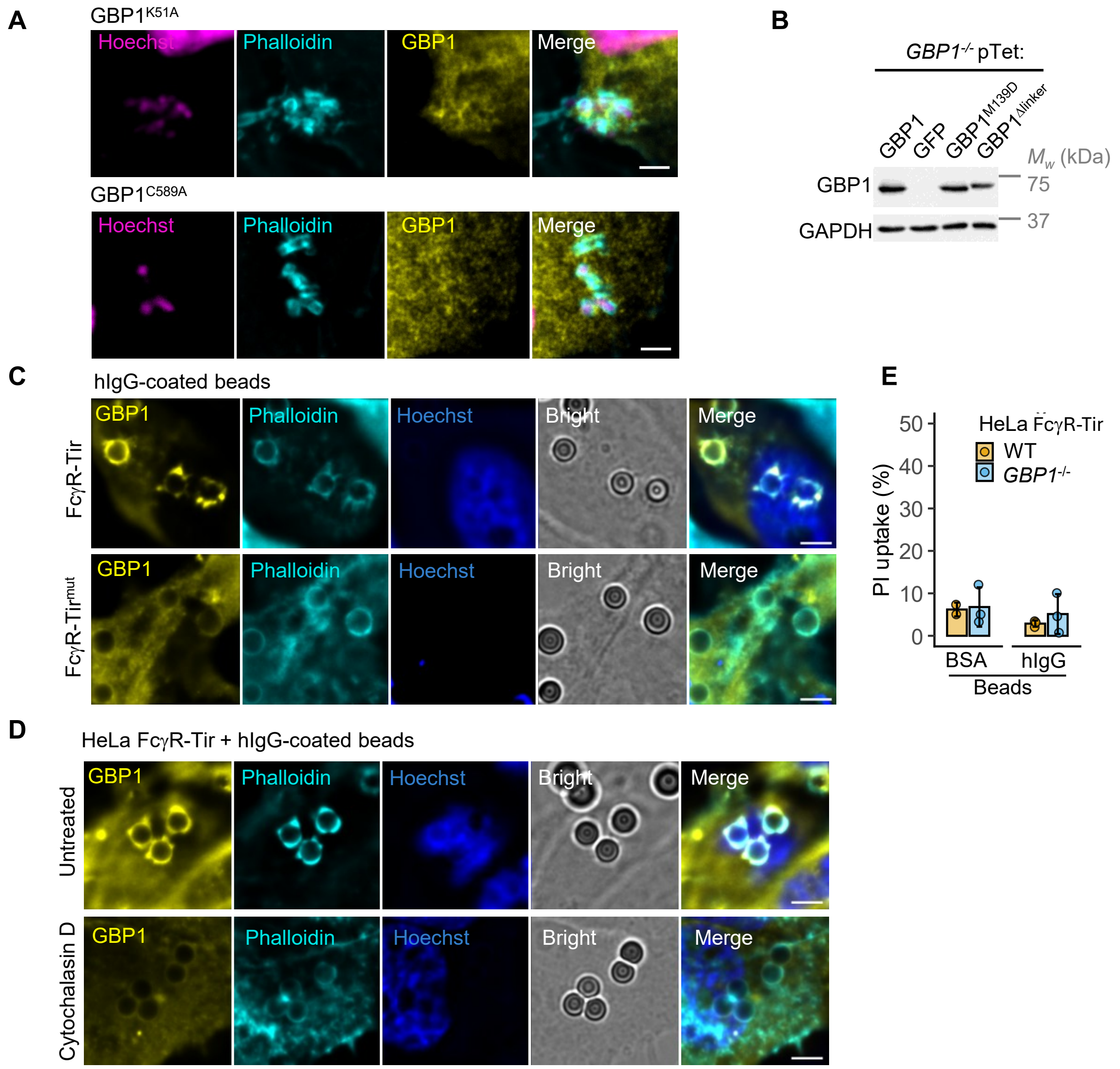


**Figure EV5. Inhibition of actin polymerisation by FcγR-Tir fusion protein blocks GBP1 recruitment to beads.**

(A) Representative images from IFNγ-primed *GBP1*$^{-/-}$ expressing GFP-tagged GBP1 K51A or C589A as indicated and infected with EPEC for 3 h. Cells were stained with anti-GBP1, phalloidin (to stain actin) and Hoechst (DNA dye for nuclei and EPEC; magenta). Scale bar, 4 µm. Data from n= 3 independent experiments.

(B) Representative immunoblots of the indicated tetracycline-controlled indicated WT or GBP1 variants in *GBP1*$^{-/-}$ cells. Cells were stably transduced with expression plasmids for the indicated proteins, treated with IFNγ (10 ng.mL$^{-1}$) and doxycycline (200 ng.mL$^{-1}$) for 16 h and cell lysates prepared for western blots. Data are representative of n = 2 independent experiments.

(C) Representative images from HeLa cells expressing FcγR-Tir or FcγR-Tir$^{mut}$ (with C-terminal mCherry2 tag) treated for 3 h with sterile polystyrene beads coated with hIg. Cells were stained with an anti-GBP1 antibody, phalloidin (to stain actin) and Hoechst (DNA dye; blue). Data are presentative of n = 3 independent experiments. Scale bar, 5 µm.

(D) Representative images from HeLa cells expressing FcγR-Tir (with C-terminal mCherry2 tag) treated for 3 h with sterile polystyrene beads coated with hIg in the absence or presence of the actin polymerisation inhibitor cytochalasin D (100 nM). Cells were stained with an anti-GBP1 antibody, phalloidin (to stain actin) and Hoechst (DNA dye; blue). Data are presentative of n = 3 independent experiments. Scale bar, 5 µm.

(E) Percentage pyroptotic cell death as measured by propidium iodide dye uptake assays of IFNγ-primed HeLa cells of the indicated genotypes stably expressing FcγR-Tir cells treated for 6 h with BSA or hIgG-coated beads as indicated. Mean ± SD error bars with symbols representing data from n = 5 independent experiments are shown.
